# The SKA complex is a mitotic vulnerability of chromosomally unstable cancers

**DOI:** 10.64898/2026.05.21.726809

**Authors:** Maud Schoot Uiterkamp, Elsa S. Coolen, Klaas de Lint, Adi Tarrab, Kruno Vukušić, Davy Rockx, Martin A. Rooimans, Tess de Kleijn, Iris J. Harmsen, Rodrigo Leite de Oliveira, Floris Foijer, René H. Medema, Iva M. Tolić, Uri Ben-David, Rob M.F. Wolthuis, Gerben Vader

**Author notes:** Department of Clinical Sciences, Faculty of Veterinary Medicine, Utrecht University; Utrecht, The Netherlands (Tess de Kleijn). These authors contributed equally to this work; author order is alphabetical (Elsa S. Coolen, Klaas de Lint, Adi Tarrab, Kruno Vukušić).

## Abstract

Aneuploidy and chromosomal instability are common characteristics of cancer but remain underexploited therapeutically. Because chromosomally unstable cells exhibit altered mitotic control, they are selectively vulnerable to inhibition of the kinesin KIF18A. Here, we identify the spindle and kinetochore–associated (SKA) complex as a mitotic regulator that establishes a similar dependency. SKA loss is tolerated in cells with a defective mitotic checkpoint, but when mitosis is prolonged, it leads to mitotic errors and loss of viability. Genetic analysis reveals that SKA and KIF18A are epistatic, although they function through mechanistically distinct mitotic pathways. SKA depletion selectively impacts chromosomally unstable cancer cells and mirrors the effects of KIF18A inhibition. We thus identify a cellular vulnerability associated with prolonged mitosis in chromosomally unstable cancers.

## Main Text

Hundreds of proteins coordinate chromosome segregation, rendering them essential for cellular survival (*e.g.,* ^1–4^). In many cases, this essentiality reflects an inherent requirement for stable genome propagation. Alternatively, lethality associated with protein loss could arise primarily from activation of the spindle assembly checkpoint (SAC) leading to prolonged mitotic arrest and cell death. The SAC, or mitotic checkpoint, senses chromosome segregation defects and halts cell cycle progression by counteracting the E3 ubiquitin ligase anaphase-promoting complex/cyclosome (APC/C) (Fig. S1A) ^5–7^. The balance between the antagonizing activities of the SAC and APC/C can therefore impact how cells respond to chromosome segregation defects. For example, inhibition or loss of the mitotic kinesin *KIF18A* activates the SAC and leads to cell death in wild-type cells, whereas cells lacking a functional SAC are resistant to KIF18A inhibition (*i.e.*, *KIF18A* is ‘synthetic viable’ in SAC-deficient conditions) ^7,8^. Many aneuploid cancers have rewired control of mitosis ^5–8^. Although this may be expected to weaken mitotic checkpoint stringency, in practice these cells often spend longer in mitosis. This characteristic, together with imbalanced chromosome numbers present in these cells, is a major determinant of KIF18A dependency ^8–11^. Based on this rationale, small-molecule inhibitors of KIF18A are in phase 1/2 clinical trials for cancers with high levels of aneuploidy/CIN ^12–15^.

### Genetic identification of SAC-contingent mitotic factors

Given the connection between SAC function and efficacy of KIF18A inhibitors, we reasoned that mapping factors whose essentiality is equally contingent on SAC function may lead to the identification of novel cancer-specific mitotic vulnerabilities. To pursue this goal, we sought to create human cells that undergo mitosis without a functional SAC. The AAA+ ATPase TRIP13 is a key component of SAC signaling ^16–18^. TRIP13 catalyzes the conformational conversion of ‘closed’ MAD2 (c-MAD2) to ‘open’ MAD2 (o-MAD2), thereby replenishing the o-MAD2 pool required for mitotic checkpoint complex (MCC) formation and SAC function (Fig. S1A) ^16–33^. *TRIP13* inactivation in near-diploid RPE1-Δ*TP53* cells (hereafter Δ*TRIP13*) altered the levels of MAD2 and the TRIP13-adaptor P31^COMET^ ^32–34^ (Fig. S1B). As expected, Δ*TRIP13* cells were unable to mount a SAC-dependent mitotic arrest after treatment with the microtubule drug nocodazole ^32–34^ (Fig. 1A, Fig. S1C). Both phenotypes were reversed upon re-expression of *TRIP13* (Fig. 1A, Fig. S1B and S1C). Despite defective checkpoint signaling, Δ*TRIP13* cells remained sensitive to *MAD2L1* loss (Fig. S1D and S1E), indicating that the ability to form an initial pool of MAD2-driven MCC remains essential ^17,32,33^. No growth defect was observed in Δ*TRIP13* cells and live-cell imaging did not reveal major defects in mitotic timing, chromosome alignment, or segregation fidelity (Fig. S1F, Movie S1 and S2, Fig. S1G). Consistently, *TRIP13* loss did not lead to significant karyotype alterations (Fig. 1B and Fig. S1H). Thus, although Δ*TRIP13* cells fail to elicit a SAC response, cell division is otherwise indistinguishable from that in wild-type cells.

**Fig. 1.**
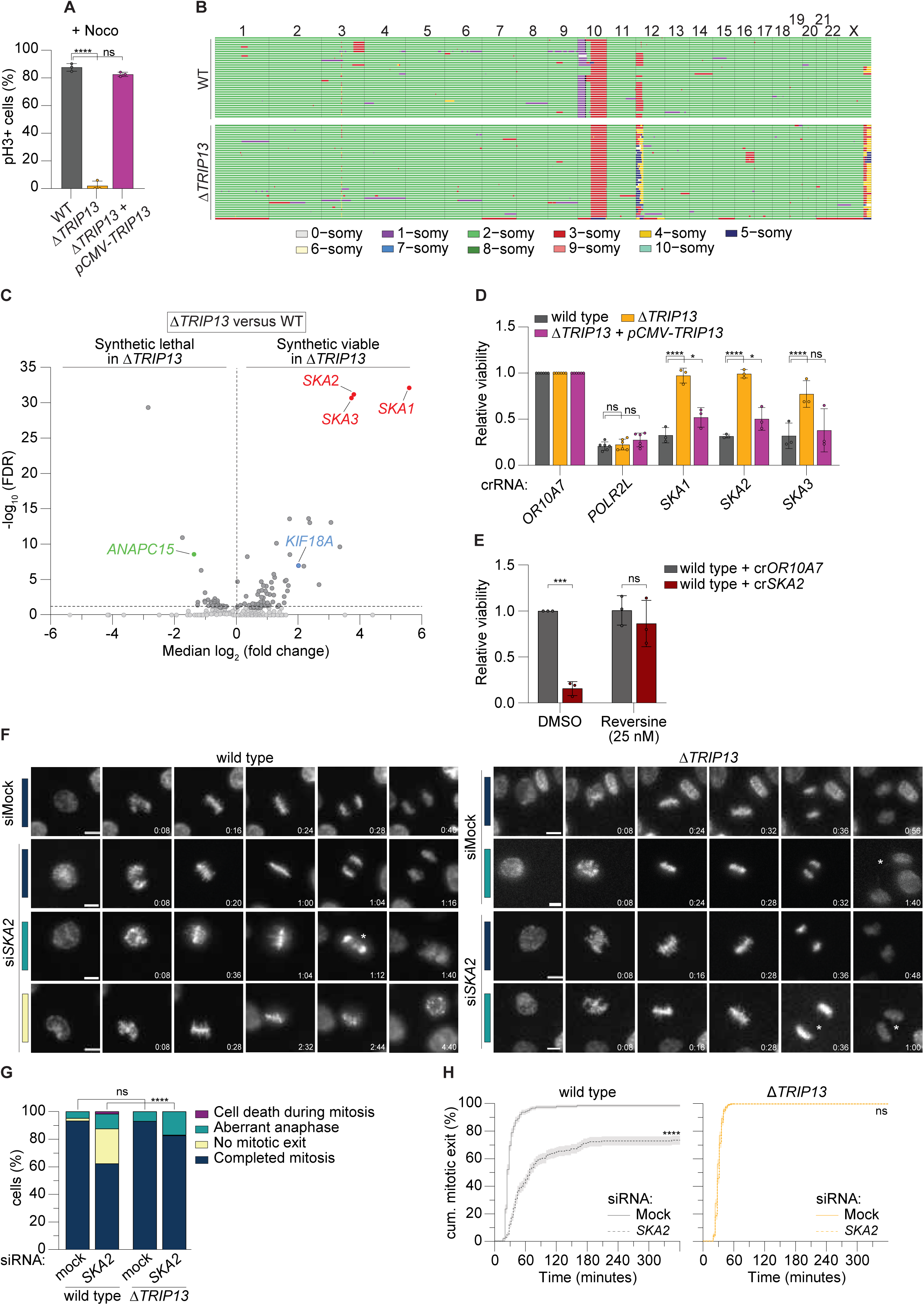
Mitotic checkpoint-dependency screen identifies the SKA complex. (**A**) Percentage of pH3-positive mitotic cells measured by flow cytometry in RPE1-*hTERT*-iCas9 Δ*TP53* (WT), RPE1-*hTERT*-iCas9 Δ*TRIP13*Δ*TP53* (Δ*TRIP13*), and RPE1-*hTERT*-iCas9 Δ*TRIP13*Δ*TP53 + pCMV-TRIP13* (Δ*TRIP13 +* p*CMV-TRIP13*) cells following 18-hour nocodazole (Noco) treatment. (**B**) Genome-wide single-cell copy number profiles of WT and Δ*TRIP13* cells (n = 48 cells). Rows represent individual cells, columns represent chromosomes. Copy number states are color-coded as indicated. Aneuploidy and karyotype heterogeneity scores were 0.079 and 0.043 for WT cells and 0.096 and 0.055 for Δ*TRIP13* cells. (**C**) Volcano plot showing genes whose loss causes synthetic lethality (SL) or synthetic viability (SV) in Δ*TRIP13* cells relative to WT cells in the CRISPR screen. Significant hits (FDR < 0.1) are shown in dark gray. (**D**) Relative viability upon CRISPR-based gene knockout. (**E**) Relative viability after CRISPR-based gene knockout and treatment with 25 nM Reversine. (**F**) Representative images of WT and Δ*TRIP13* cells treated with mock or *SKA2* siRNA and stained with a live-cell DNA probe, undergoing division. Cell fates are color-coded: blue bars indicate completed mitosis, yellow bars indicate lack of mitotic exit during the imaging period, teal bars indicate an aberrant anaphase, purple bars indicate mitotic cell death. An asterisk is placed to identify aberrancies. Time is indicated in hours:minutes. Scale bar, 10 μm. (**G**) Quantification of mitotic cell fates of WT and Δ*TRIP13* cells by live-cell imaging after treatment with mock or *SKA2* siRNA. For (F) and (G), 169, 178, 221, and 195 cells were analyzed for WT siMock, WT siSKA2, Δ*TRIP13* siMock, and Δ*TRIP13* siSKA2 cells, respectively. (**H**) Mitotic exit timing of WT and Δ*TRIP13* cells after mock or *SKA2* siRNA treatment. Cells were tracked for more than 20 hours, but no new mitotic exits were observed after 6 hours. Only cells observed entering mitosis were included in this analysis. Shaded area indicates standard error. Data in (A), (D), (E), (G), and (H) were obtained from three biologically independent experiments, with technical duplicates in (A) and triplicates in (D-E). Statistical significance was assessed by one-way ANOVA (A), two-way ANOVA (D-E), Fisher’s exact test (G), or log-rank test (H), each with correction for multiple comparisons. *P < 0.05; ***P < 0.001; ****P < 0.0001; ns, not significant.

To comprehensively identify factors whose essentiality is impacted by SAC functionality, we performed a genome-wide CRISPR-Cas9 knockout screen in Δ*TRIP13* cells (Fig. S1I). This identified two classes of genetic interactions. In one class, genes became more essential when *TRIP13* was absent (‘synthetic lethality’ (SL)). In the other, genes became less essential in Δ*TRIP13* cells (‘synthetic viability’ (SV)). We identified 74 SL and 46 SV genes (Fig. 1C, data S1A-C). The APC/C subunit APC15 (encoded by *ANAPC15*) scored as a strong SL hit, validating the screen (Fig. 1C) ^17^. Indeed, combined loss of *TRIP13* and *ANAPC15* triggered a mitotic (metaphase) arrest, attributed to an inability to disassemble the MCC ^17,35–37^, resulting in viability loss (Fig. S1J-M).

Inspired by the reported synthetic viable interaction between KIF18A and the mitotic checkpoint ^7^, we focused our attention on the identified SV genes. Gene ontology (GO) term enrichment analysis of the 46 SV genes revealed significant enrichment for factors involved in mitotic spindle dynamics and chromosome segregation (Fig. S2A). Gene Effect Scores for these SV genes (*i.e*., ‘essentiality’ scores), derived from the DepMap initiative ^38^, skewed towards values typical of genes that are classified as ‘common essential’ (Fig. S2B). Taken together, these data hint that these SV genes become essential mainly because their loss triggers a checkpoint-dependent mitotic arrest.

### Essentiality of the SKA complex in mitosis is determined by the mitotic checkpoint

Notably, the top SV hits encode all subunits of the spindle and kinetochore–associated (SKA) complex (SKAc): *SKA1, SKA2*, and *SKA3* (Fig. 1C). This trimeric complex stabilizes kinetochore-microtubule attachments and is essential for anaphase onset and mitotic exit ^39–45^. Two explanations could account for the observed SV behavior: *i*) disruption of the identified gene could cause viability loss in wild-type cells, which is suppressed in Δ*TRIP13* cells, or *ii*) gene disruption could rescue a proliferation defect present in Δ*TRIP13* cells. Analysis of SKAc gRNA abundance demonstrated behavior that is compatible with the former scenario (Fig. S2C). Viability and clonogenic assays validated SV behavior upon *SKA1*, *SKA2*, or *SKA3* loss (Fig. 1D and S2D). By using the small-molecule inhibitor Reversine against the essential SAC component MPS1 ^46^ (Fig. S1A), we confirmed that synthetic viability of the SKAc in Δ*TRIP13* cells was driven by broad SAC dysfunction (Fig. 1E and Fig. S2E). Our screen thus identified a selected list of mitotic factors, prominently among them *SKA1*-*3*, whose viability defects are mitigated by SAC loss.

We tracked the first mitosis after siRNA-mediated knockdown of *SKA2* by live-cell imaging (experimental set-up in Fig. S3A and S3B). In wild-type cells, *SKA2* knockdown increased the fraction of cells in metaphase from 2% to 25% (Fig. 1F and 1G, Fig. S3C, Movie S3-6) and doubled mitotic duration from 30±1 to 65±4 minutes (mean±SEM, Fig. 1H, Fig. S3D), consistent with prior reports ^39–45^. The fraction of aberrant anaphases increased from 5% to 11%, and chromosome alignment was delayed (Fig. 1F and 1G, Fig. S3C and S3E). Conversely, *SKA2* depletion did not induce a metaphase arrest in Δ*TRIP13* cells (Fig. 1F-H, Fig. S3C, Movie S7-10). Instead, these cells progressed through mitosis unimpeded within 31±1 minutes (mean±SEM) (Fig. S3D). Overall, these results demonstrate that loss of TRIP13 circumvents the metaphase arrest induced by *SKA2* depletion, explaining the synthetic viability observed in Δ*TRIP13* cells. We also created stable *SKA1, SKA2,* or *SKA3* knockouts in the Δ*TRIP13* background. Consistent with prior studies, loss of any of the three SKA subunits destabilized the entire SKA complex (Fig. S3F) ^40,41^. Re-expression of *TRIP13* resulted in a near-complete loss of viability in these cells (Fig. S3G), indicating that no compensatory, adaptive mechanisms were selected. Mitotic phenotyping of these double knockouts revealed that approximately 86% of all cells progressed through mitosis efficiently; 12% of cells underwent aberrant anaphase compared to 3% of wild-type or Δ*TRIP13* cells (Fig. S4A and S4B, Movie S11-16). Time to metaphase alignment and total mitotic duration remained similar, with a mitotic duration of 32±1 minutes in Δ*TRIP13*, 34±1 minutes in Δ*SKA2*Δ*TRIP13,* and 36±1 minutes in Δ*SKA3*Δ*TRIP13* cells (mean±SEM, Fig. S4C-E). Consistent with efficient and robust mitotic progression, chromosome counts of Δ*SKA2*Δ*TRIP13* cells were not significantly different from that of either wild-type or Δ*TRIP13* cells, with a mean chromosome number per cell of 45.8±0.1 in wild-type cells, 46.5±1.5 in Δ*TRIP13*, and 45.3±1.5 in Δ*SKA2*Δ*TRIP13* (mean±SEM, Fig. S1H). These findings indicate that SKAc loss impacts viability by causing an extended mitotic arrest in cells with an intact mitotic checkpoint.

We next asked what molecular defect underlies the lethality of SKAc loss in checkpoint-proficient cells. One possibility is that unstable kinetochore-microtubule attachments may repeatedly re-trigger the SAC ^47–55^. Another possibility is that the SKAc has a direct role in promoting mitotic exit, as has been suggested ^39,43,56^. In agreement with a defect that originates from suboptimal kinetochore-microtubule interactions, we found that Δ*SKA2*Δ*TRIP13* cells showed a trend of increased sensitivity to the microtubule-destabilizer Noscapine (Fig. S4F) ^57^.

Like the SKAc, *KIF18A* emerged as an SV interactor of *TRIP13* (Fig. 1C), as expected based on earlier reports ^7^. Indeed, Δ*TRIP13* cells were less sensitive to *KIF18A* knockout or chemical inhibition using Sovilnesib as compared to wild-type cells ^58^ (Fig. S4G). In addition to *SKA1-3* and *KIF18A*, our screen identified several additional mitotic regulators. These included multiple subunits of the Augmin complex, which promotes microtubule-dependent microtubule nucleation ^59–66^, as well as the protein kinase *CSNK1A1*, the E3 ligase subunit *CUL3*, the spliceosome component *MAGOH*, and the phosphatase complex *PPP1CB-PPP1R12A* (Fig. 2A, data S1B) ^67–73^. All showed SAC-dependent effects on viability (Fig. S5A-C). In contrast, core mitotic regulators such as *PLK1*, *AURKA*, *NDC80* and *NUF2* remained essential regardless of *TRIP13* status (Fig. S5D). Thus, our SV hits form a unique subset of mitotic regulators defined by SAC-contingent essentiality.

**Fig. 2.**
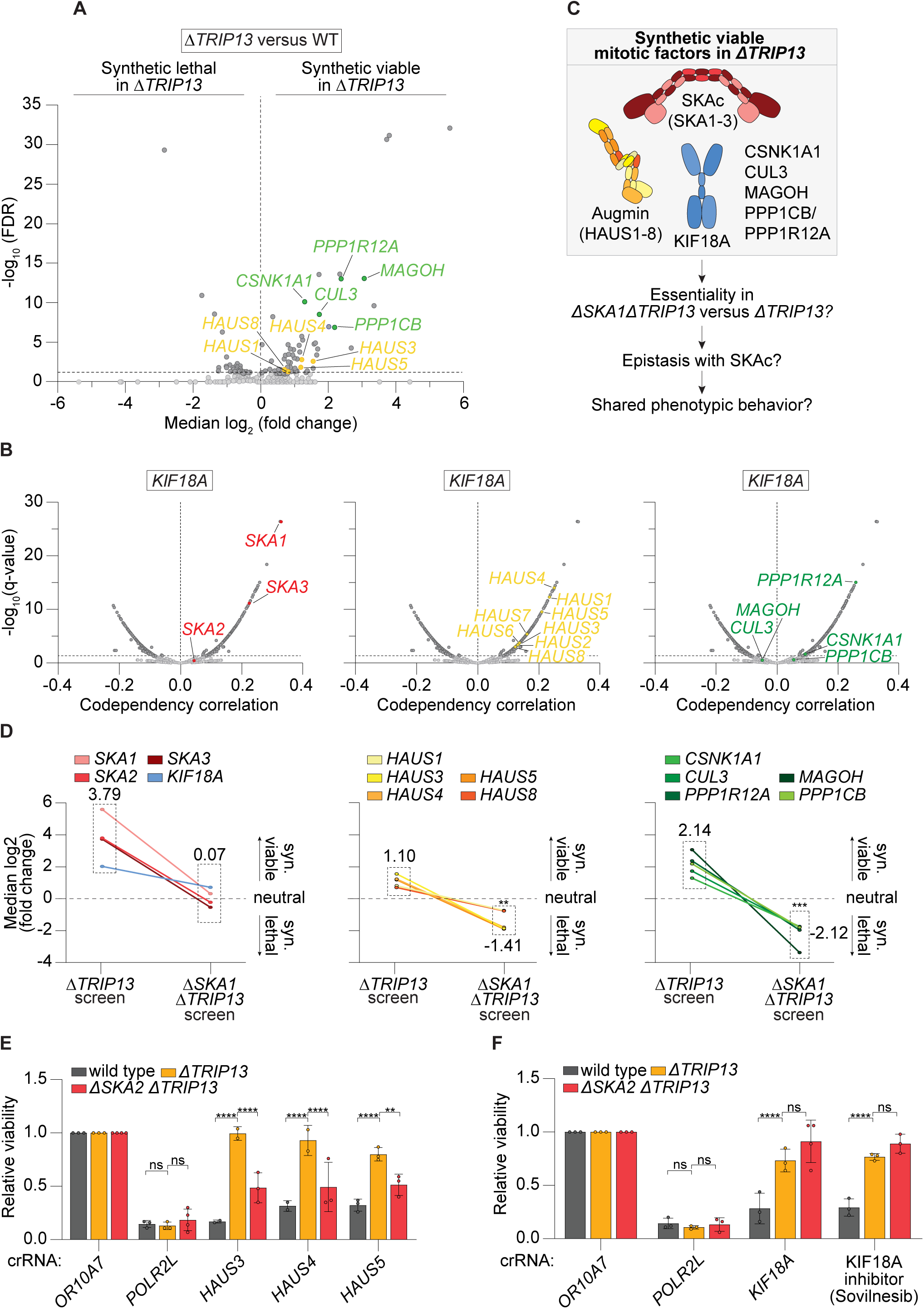
Co-dependency and genetic epistasis between the SKA complex and KIF18A. (**A**) Volcano plot showing mitosis-linked genes whose loss causes synthetic viability (SV) in RPE1-*hTERT*-iCas9 Δ*TRIP13*Δ*TP53* (Δ*TRIP13*) cells relative to RPE1-*hTERT*-iCas9 Δ*TP53* (WT) cells in the CRISPR screen. Significant hits (FDR < 0.1) are shown in dark gray. (**B**) Volcano plots showing gene co-dependency correlations with *KIF18A* across cancer cell lines. The x-axis represents the Pearson correlation coefficient (r), the y-axis shows the significance. Genes with significant co-dependency (q < 0.05) are shown in dark gray. (**C**) Schematic depicting SAC-contingent synthetic viable factors and the rationale for the Δ*SKA1*Δ*TRIP13* versus Δ*TRIP13* setup to derive epistatic relationships. (**D**) Comparison of gene effects in the Δ*TRIP13* and Δ*SKA1*Δ*TRIP13* CRISPR screens based on the mean abundance scores of four sgRNAs per gene. Plotted genes were significant synthetic viable hits in the Δ*TRIP13* screen (positive fold change in guide abundance). Significance is assessed through one-way ANOVA with correction for multiple comparisons, by comparison to the mean of *SKA1-3* and *KIF18A* guide abundances in the Δ*SKA1*Δ*TRIP13* screen. (**E** and **F**) Relative viability upon CRISPR-based gene knockout or chemical inhibition. n = 2-4 biologically independent experiments with technical triplicates. Two-way ANOVA with correction for multiple comparisons, **P < 0.01; ****P < 0.0001; ns, not significant.

### Genetic co-dependence between KIF18A and SV factors

To explore whether the genetic parallels between *KIF18A* and SV factors observed in SAC-deficient RPE1 cells can be extrapolated to other (SAC proficient) cellular contexts, we analyzed CRISPR-mediated loss-of-function effects in a large cohort of human cancer cell lines, available from the DepMap initiative ^38^. Comparing the impact on cellular survival of loss of SV factors to that of KIF18A interestingly revealed significant genetic similarities (*i.e.*, co-dependence) between *KIF18A* and *SKA1, SKA3, HAUS1-8*, *CSNK1A1* and *PPP1R12A* across a wide range of cancer cell lines (Fig. 2B). We note that *SKA1* exhibited the second highest *KIF18A* co-dependency score of all genes (∼18,000) investigated in the DepMap initiative. Reciprocal co-dependency analyses using *SKA1-3* and *HAUS1-8* as query genes revealed similar relationships (Fig. S6A-S6C). These results recapitulate the differential viability effects observed in our SAC-deficiency screen. As mitotic checkpoint-dependent essentiality underlies the selective killing of chromosomally unstable cancer cells by KIF18A inhibition ^7,8^, this analysis implies that targeting such SV factors may expose comparable cancer-specific vulnerabilities.

### KIF18A and the SKA complex are epistatic

To position the identified SAC-contingent SV factors in functional pathways relative to our top hits *SKA1-3*, we performed a second genetic screen in *ΔSKA1ΔTRIP13* cells (Fig. S7A, data S1D). Comparing gene-loss effects in *ΔTRIP13* versus *ΔSKA1ΔTRIP13* cells allowed us to identify factors that genetically phenocopy or diverge from SKA complex loss alone (Fig. 2C). This can reveal genetic relationships with the SKAc, which may indicate shared phenotypic behavior. As expected from components of the same protein complex, loss of *SKA2* or *SKA3* had no differential effect in Δ*SKA1*Δ*TRIP13* cells (*i.e.*, their gene-loss effect became neutral, similar to the effect of targeting *SKA1*) (Fig. 2D). Most other SV factors behaved differently. Augmin complex components (*HAUS1, 3-5* and *8)*, as well as *CSNK1A1, CUL3, MAGOH, PPP1R12A,* and *PPP1CB* shifted from synthetic viable in Δ*TRIP13* cells to synthetic lethal in Δ*SKA1*Δ*TRIP13* cells (Fig. 2D). These findings indicate that, contrary to SKA1-3 loss, disruption of these factors causes additional – presumably mitotic – defects that negatively impact cellular viability in the absence of the SKAc. Unlike all other mitotic SV genes, KIF18A loss had no additional effect on viability once the SKAc was absent (*i.e.*, KIF18A loss became neutral in Δ*SKA1*Δ*TRIP13* cells, mirroring the behavior of *SKA2* and *SKA3* (Fig. 2D)). In genetic terms, these data point to a unique epistatic relationship between *KIF18A* and *SKA1-3*.

Confirming these analyses, viability of Δ*SKA*2Δ*TRIP13* or Δ*KIF18A*Δ*TRIP13* cells was decreased upon concurrent loss of Augmin subunits, *CSNK1A1, CUL3, MAGOH, PPP1R12A,* or *PPP1CB* (Fig. 2E, Fig. S7B). In contrast, loss or chemical inhibition of *KIF18A* did not reduce cellular survival in Δ*SKA*2Δ*TRIP13* cells (Fig. 2F). Similarly, removal of *SKA2* in Δ*KIF18A*Δ*TRIP13* cells (Fig. S7C) did not impact viability (Fig. S7D). We conclude that, among the larger group of SAC-contingent SV factors, the SKA complex and KIF18A exhibit an epistatic relationship, indicating that the SKAc and KIF18A impact cellular survival in comparable ways.

### The SKA complex is a vulnerability in chromosomally unstable cancer cells

Cancers that are aneuploid and/or exhibit chromosomal instability (CIN) specifically depend on KIF18A during mitosis, and the use of small-molecule inhibitors of KIF18A is currently being explored to combat such cancers ^8–15^. The identified genetic similarities between the SKA complex and KIF18A inspired us to focus on the SKA complex and investigate whether it represents a similar vulnerability in chromosomally unstable cancers. As large-scale dependency datasets do not typically capture ongoing chromosomal instability, we used aneuploidy status as a related feature to explore this possibility. The association between aneuploidy and increased sensitivity to KIF18A loss has previously been established by dependency analysis in a large cohort of cancer cell lines ^9–11^. We therefore analyzed genome-wide CRISPR dependency data from the CCLE ^38^. Interestingly, stratification by aneuploidy status showed that the top 25% most aneuploid cancer cell lines were significantly more dependent on *SKA1* than the 25% most euploid cancer cell lines, mirroring their increased dependency on KIF18A (Fig. 3A) ^9^. Similar aneuploidy-associated dependence was observed for the complete SKAc (*SKA1-3*) (Fig. 3A). Across the screened cancer cell lines, higher aneuploidy scores were modestly but significantly correlated with increased dependency on *SKA1* and *SKA3,* reflected by more negative dependency scores (Fig. S8A). Again, these effects were comparable to those seen for *KIF18A* (Fig. S8B). Most other SV hits showed no association with aneuploidy scores (Fig. S8C and S8D). These data show that, indeed, loss of the SKAc exposes a KIF18A-like, aneuploidy-specific cancer cell vulnerability across a large cohort of cancer cell lines.

**Fig. 3.**
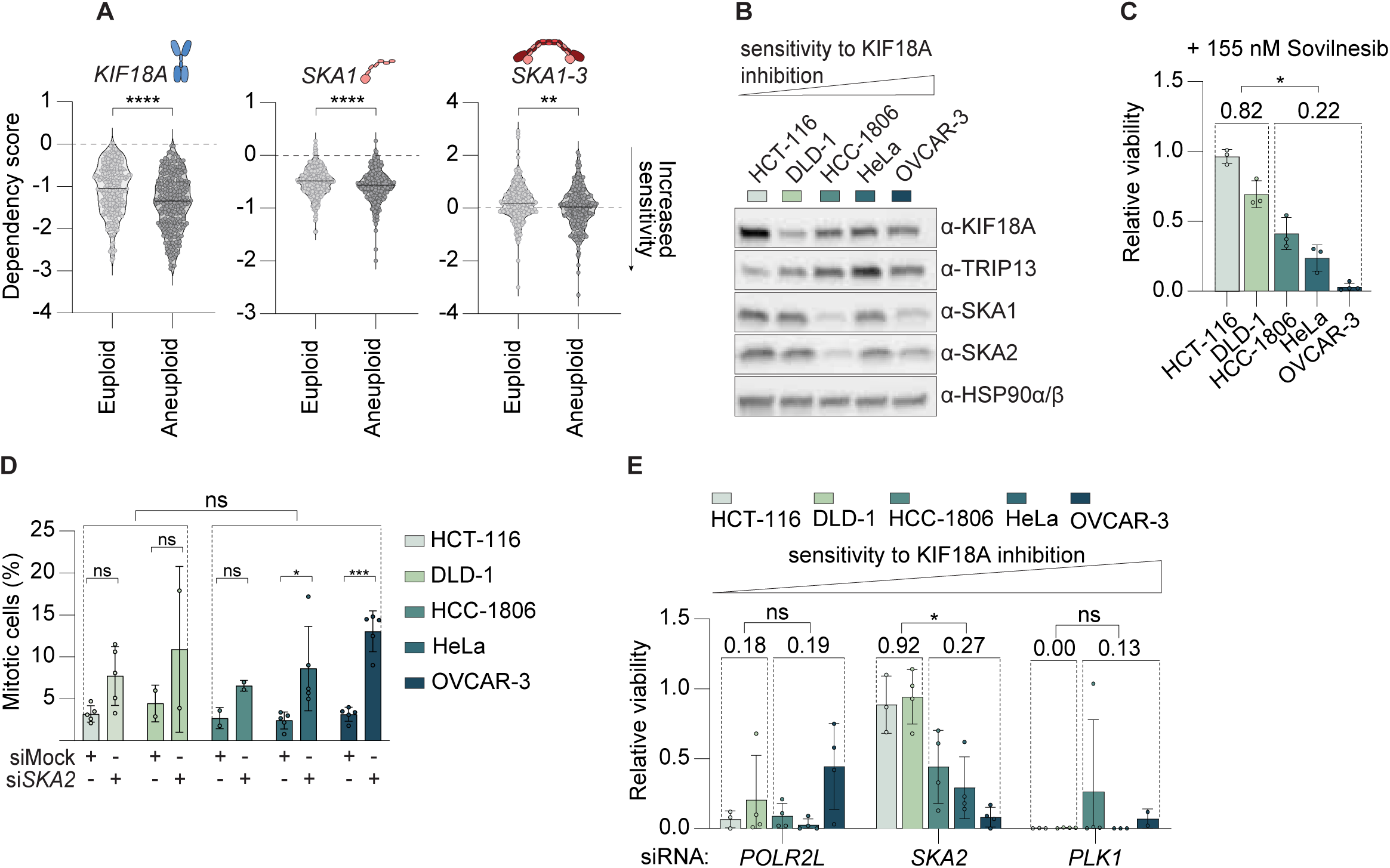
The SKAc is a cellular vulnerability in chromosomally unstable cancer cells. (**A**) Sensitivity of near-euploid and highly aneuploid cancer cell lines to loss of *KIF18A* (left), *SKA1* (middle), or *SKA1-3* (right), as measured in the Chronos CRISPR-Cas9 dataset. More negative dependency scores indicate greater gene essentiality. (**B**) Representative western blot of KIF18A, TRIP13, SKA1, and SKA2 across a panel of cancer cell lines with differing sensitivity to KIF18A inhibition. (**C**) Relative viability of cancer cell lines with differing sensitivity to KIF18A inhibition after treatment with 155 nM Sovilnesib, a KIF18A inhibitor. (**D**) Percentage of pH3-positive mitotic cells measured by flow cytometry after treatment with mock or *SKA2* siRNA for 60 hours. Grouped as KIF18A-inhibition-insensitive (HCT-116, DLD-1) and KIF18A-inhibition-sensitive (HCC-1806, HeLa, OVCAR-3). Statistical comparison between groups are based on mitotic fraction upon si*SKA2* treatment. (**E**) Relative viability after SKA2 depletion, grouped as KIF18A-inhibition-insensitive (HCT-116, DLD-1) and KIF18A-inhibition-sensitive (HCC-1806, HeLa, OVCAR-3). Data in (C) and (E) were obtained from three or four biologically independent experiments with technical triplicates, and data in (D) were obtained from two or five biologically independent experiments with technical duplicates. Statistical significance was assessed by two-tailed unpaired t-test (A), nested two-tailed t-test (C and E), two-way ANOVA with correction for multiple comparisons (D), and nested t-test (D). *P < 0.05; **P < 0.01; ***P = 0.0002; ****P < 0.0001; ns, not significant.

We next experimentally tested whether SKAc loss mimics KIF18A inhibition in cancer cell lines ^8,74^. To do so, we used a panel of colon (HCT-116, DLD-1), breast (HCC-1806), ovarian (OVCAR-3), and cervical (HeLa) cancer cell lines, previously shown to span a range of KIF18A inhibitor sensitivities and aneuploidy levels ^8,74^ (Fig. 3B). Based on drug response curves to the KIF18A inhibitor Sovilnesib ^58^ (Fig. 3C, Fig. S9A) that aligned with published data ^8,74^, we categorized HCT-116 and DLD-1 as insensitive cell lines, and HCC-1806, HeLa, and OVCAR-3 as sensitive. Consistent with the roles of the SKAc in mitosis ^39–45^, siRNA-mediated depletion of *SKA2* (Fig. S9B) increased the mitotic fraction in all cell lines tested (Fig. 3D), confirming efficient siRNA-mediated protein depletion. Strikingly, viability of the KIF18Ai-insensitive cell lines remained at an average of 92% upon *SKA2* depletion, while it dropped to 27% in sensitive cell lines (Fig. 3E), mirroring the response to KIF18A inhibition (Fig. 3C) ^8^. This differential sensitivity could not be attributed to different efficacies of siRNA-mediated protein depletion, as knockdown of the essential factor *POLR2L* resulted in efficient viability loss (Fig. 3E). Further, the effects were not caused by differences in baseline proliferation rates (Fig. S9C), or by different intrinsic vulnerabilities to broad mitotic disruption, as *PLK1* knockdown reduced viability to below 15% in all cell lines (Fig. 3E). The fact that we observed comparable mitotic fractions upon SKA2 depletion while viability effects differed starkly across the panel of cells lines could indicate that *i) SKA2*-sensitive cells are more prone to cell death when arrested in mitosis, and/or ii) these cells arrest for a longer time in mitosis (Fig. 3D and E).

Notably, depletion of other SV hits (*CSNK1A1, CUL3, MAGOH, PPP1R12A,* and *PPP1CB*) did not reproduce the SKA-like differential effects on viability or mitotic arrest in this cancer cell panel (Fig. S9D and S9E). This reinforces our prior epistasis analysis that isolated the SKAc as a uniquely KIF18A-like factor among a larger group of SAC-contingent mitotic genes. Taken together, we conclude that the SKA complex is a selective cellular vulnerability in chromosomally unstable cancer cells.

### The SKA complex and KIF18A differentially impact chromosome alignment in mitosis

Considering the similar impact of interference with the SKAc and KIF18A on cellular viability and mitotic defects, we investigated how these factors mechanistically relate to each other during mitosis. SKA proteins and KIF18A exhibit proximity at kinetochores during (pro)metaphase ^75,76^. SKAc localization at kinetochores was maintained in *ΔKIF18AΔTRIP13* cells, and KIF18A remained robustly associated with kinetochore microtubules in *ΔSKA2ΔTRIP13* cells (Fig. 4A). In each case, signal was absent in the corresponding knockout line, confirming specificity. These observations indicate that these proteins localize independently, supporting a model in which the SKAc and KIF18A interact functionally through kinetochore–microtubule mechanics rather than via direct, reciprocal recruitment.

**Fig. 4.**
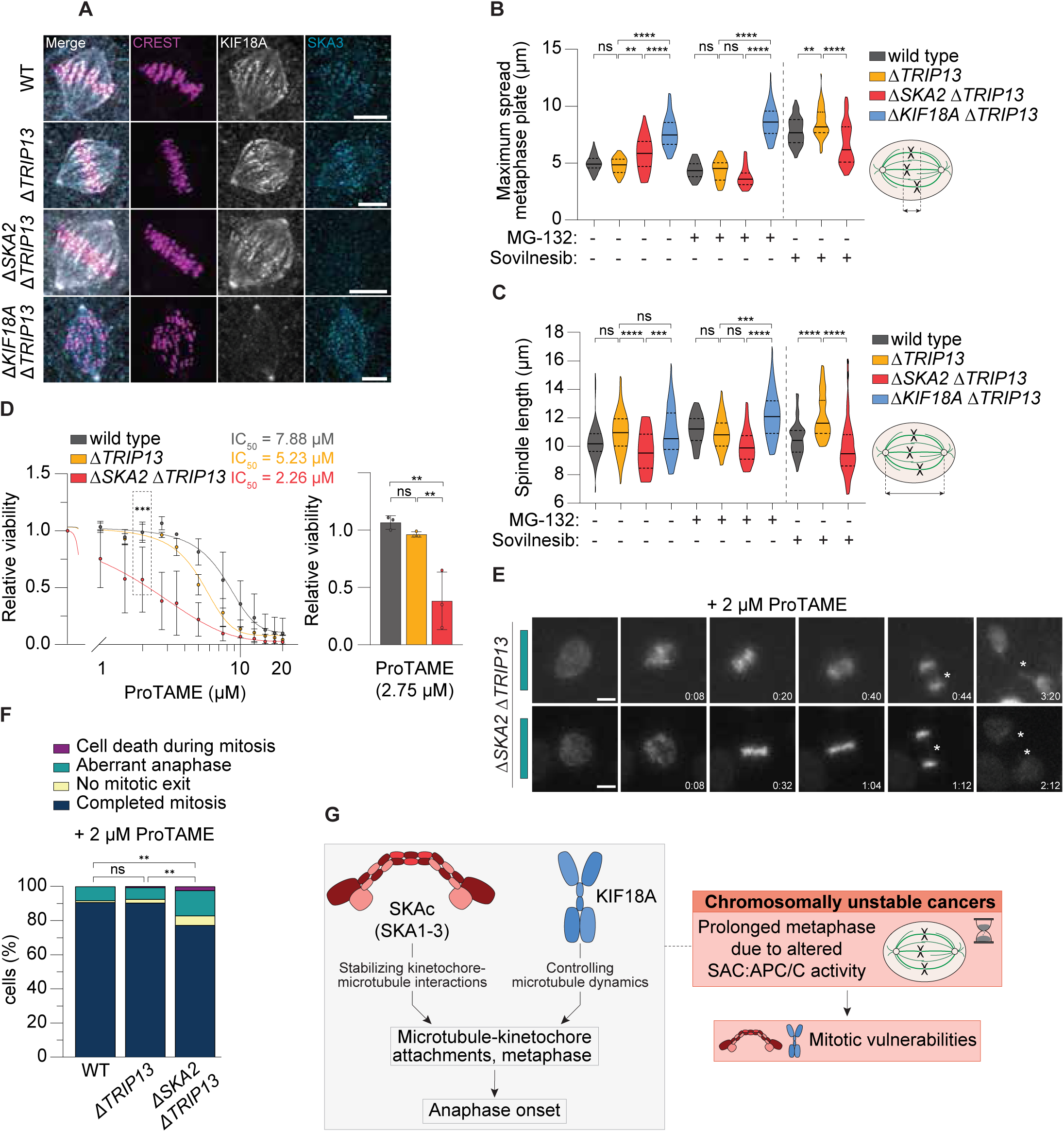
Differential impact of SKA complex and KIF18A on cell division, with SKA-deficient defects exacerbated by prolonged mitosis. **(A**) Representative images of RPE1-*hTERT*-iCas9 Δ*TP53* (WT), RPE1-*hTERT*-iCas9 Δ*TRIP13*Δ*TP53* (Δ*TRIP13*), RPE1-*hTERT*-iCas9 Δ*SKA2*Δ*TRIP13*Δ*TP53* (Δ*SKA2*Δ*TRIP13*), and RPE1-*hTERT*-iCas9 Δ*KIF18A*Δ*TRIP13*Δ*TP53* (Δ*KIF18A*Δ*TRIP13*) cells stained for CREST (magenta), KIF18A (black), and SKA3 (cyan). Scale bar, 5 μm. (**B**) Spindle length of WT, *ΔTRIP13*, *ΔSKA2ΔTRIP13*, and *ΔKIF18AΔTRIP13* cells, with or without MG-132 or Sovilnesib treatment. For WT, Δ*TRIP13*, Δ*SKA2*Δ*TRIP13*, and Δ*KIF18A*Δ*TRIP13*, 49, 54, 40, and 47 cells were analyzed, respectively. For MG-132-treated WT, Δ*TRIP13*, Δ*SKA2*Δ*TRIP13*, and Δ*KIF18A*Δ*TRIP13*, 45, 30, 38, and 46 cells were analyzed, respectively. For Sovilnesib-treated WT, Δ*TRIP13*, and Δ*SKA2*Δ*TRIP13*, 49, 47 and 36 cells were analyzed, respectively. (**C**) Maximum metaphase plate spread of WT, *ΔTRIP13*, *ΔSKA2ΔTRIP13*, and *ΔKIF18AΔTRIP13* cells, with or without MG-132 or Sovilnesib treatment. For WT, Δ*TRIP13*, Δ*SKA2*Δ*TRIP13*, and Δ*KIF18A*Δ*TRIP13*, 49, 53, 40, and 48 cells were analyzed, respectively. For MG-132-treated WT, Δ*TRIP13*, Δ*SKA2*Δ*TRIP13*, and Δ*KIF18A*Δ*TRIP13*, 45, 30, 38, and 46 cells were analyzed, respectively. For Sovilnesib-treated WT, Δ*TRIP13*, and Δ*SKA2*Δ*TRIP13*, 52, 49, and 36 cells were analyzed, respectively. (**D**) Dose-response curve of relative viability after treatment with varying concentrations of ProTAME, and relative viability upon 2.75 μM of ProTAME. The concentration of 2 μM, indicated with the dashed lines, was used for subsequent experiments. (**E**) Representative images of Δ*SKA2*Δ*TRIP13* cells stained with a live-cell DNA probe, undergoing division after treatment with 2 μM ProTAME. Teal bars indicate aberrant anaphase. An asterisk is placed to identify aberrancies. Time is indicated in hours:minutes. Scale bar, 10 μm. (**F**) Quantification of mitotic cell fates by live-cell imaging after treatment with 2 µM ProTAME. 194, 281, and 186 cells were analyzed for WT, Δ*TRIP13*, and Δ*SKA2*Δ*TRIP13* cells, respectively. (**G**) A model depicting the identification of the SKAc as a KIF18A-like mitotic vulnerability of chromosomally unstable cancers, influenced by mitotic duration. SKAc and KIF18A impact chromosome alignment through distinct mechanisms. Data in (B) and (C) were obtained from three or four biologically independent experiments, data in (D-F) were obtained from three biologically independent experiments, with technical triplicates in (D). Statistical significance was assessed by two-way ANOVA with correction for multiple comparisons (B-D), and pairwise Fisher’s exact test with correction for multiple comparisons (F). *P < 0.05; **P < 0.01; ***P < 0.001; ****P < 0.0001; ns, not significant.

To understand why the SKAc and KIF18A behave similarly genetically, we compared their mitotic phenotypes. Consistent with its established roles in suppressing chromosome oscillations, promoting chromosome alignment, and suppressing microtubule growth ^7,76^, loss of KIF18A significantly increased chromosome spread in metaphase (measured as metaphase plate width and full width at half maximum of the metaphase plate) in Δ*TRIP13* cells (Fig. 4B, Fig. S10 and S10B). Prolonging mitosis by a 30-minute metaphase arrest with the proteasome inhibitor MG-132 did not rescue this chromosome alignment defect (Fig. 4B, Fig. S10B). KIF18A loss also significantly increased spindle length (Fig. 4C, Fig. S10B). In contrast, loss of SKA2 had only modest effects on chromosome alignment, which were largely normalized following mitotic prolongation with MG-132 (Fig. 4B, Fig. S10B). Loss of SKA2 also reduced the average spindle length during unperturbed mitosis (Fig. 4C, Fig. S10B). Together, these results show that SKAc loss does not completely phenocopy KIF18A loss. Notably, deletion of *SKA2* in the *ΔTRIP13* background markedly reduced the severe chromosome alignment defects caused by acute KIF18A inhibition (Fig. 4B, Fig. S10B). This suppression occurred independently of mitotic prolongation. Thus, although SKAc and KIF18A loss have similar effects on cell viability, they do not disrupt mitosis in the same way. Rather, our data indicate that the SKA complex is required for the full manifestation of the chromosome alignment defects caused by KIF18A loss. These observations reinforce the current model that while the SKA complex is needed to stabilize end-on kinetochore–microtubule attachments and to promote persistent coupling to dynamic microtubule tips ^39–45^, KIF18A is essential for microtubule dynamics, suppression of chromosome oscillations, and coordinated kinetochore–microtubule interactions ^7,76^. Taken together, our data indicate that the SKAc and KIF18A have opposing contributions to microtubule-kinetochore attachments and thus stable chromosome alignment. These observations may explain their epistatic genetic relationship.

### Prolonged mitosis exposes the SKA complex as a cellular vulnerability

Our data indicate that the mitotic SKAc function is molecularly distinct from that of KIF18A and that SKAc loss may even suppress KIF18A loss-associated defects in chromosome alignment. However, the loss of these factors exposes highly similar cellular vulnerabilities. Based on our analyses and the known roles of the SKAc, we consider it likely that the effects of SKAc loss originate from mitotic defects that are specifically exposed in chromosomally unstable cancer cells. In such cancer cells, cellular sensitivity to KIF18A inhibition is linked to molecular rewiring of mitotic control ^8^. This altered behavior manifests as prolonged metaphase duration, likely driven by the altered balance between SAC and APC/C activities. We therefore explored whether prolonging metaphase, via interference with APC/C activity using proTAME ^77^, impacted cellular viability and chromosome segregation in Δ*SKA*2Δ*TRIP13* cells. Indeed, Δ*SKA*2Δ*TRIP13* cells were significantly more sensitive to partial APC/C inhibition, with an IC_50_ shift from 5.2 μM in Δ*TRIP13* to 2.3 μM in Δ*SKA*2Δ*TRIP13* cells (Fig. 4D). Live-cell imaging following proTAME treatment (Fig. S10C) revealed a three-fold increase in mitotic cell death and more than a doubling of aberrant anaphases in Δ*SKA*2Δ*TRIP13* cells compared to Δ*TRIP13* alone upon induction of a mitotic delay (Fig. 4E and 4F, Fig. S10D, Movie S17-18). These findings indicate that while SAC deficiency permits mitotic progression despite SKA complex loss, SAC-deficient cells remain sensitive to elongated metaphases. This direct connection between SKA dependency and interference with APC/C activity mirrors behavior observed upon KIF18A inhibition ^8^. Combined, these experiments identify the SKAc as a novel vulnerability of chromosomally unstable cancer cells that is exposed in cells experiencing prolonged mitosis.

## Discussion

Using genetic perturbations, we identify a class of mitotic regulators that becomes essential only when the spindle assembly checkpoint (SAC) is intact. This finding suggests that the mitotic defects caused by loss of such factors are not intrinsically lethal but only become detrimental when checkpoint engagement prolongs mitosis. Among these factors is the kinesin KIF18A, which has recently emerged as a selective mitotic vulnerability in chromosomally unstable cancers ^8–11^.

Our analyses in both non-transformed and cancer cell models identify the SKA complex as a KIF18A-like dependency (Fig. 4G). Like KIF18A, the SKAc is dispensable when the mitotic checkpoint is disabled but required when SAC signaling is intact. Genetically, the SKAc and KIF18A behave epistatically. Consistent with this connection, analysis of CRISPR dependency data from a large cohort of cancer cell lines reveals that SKAc dependency is associated with elevated aneuploidy levels, paralleling previous observations for KIF18A ^8^. We corroborate this association experimentally, demonstrating a selective requirement for the SKAc in aneuploid cancer cells. Although the SKAc and KIF18A impact kinetochore-microtubule interactions through molecularly distinct mechanisms, we find that SKAc loss, like KIF18A inhibition/loss, triggers persistent mitotic checkpoint activation which is specifically deleterious in cells with prolonged metaphase (*8)*. Indeed, defects in chromosome segregation caused by SKAc loss are tolerated when the SAC is disrupted but become more severe as mitotic duration increases. Together, these findings suggest that the selective importance of the SKAc in chromosomally unstable cancer cells arises from chromosome segregation defects that coincide with prolonged checkpoint-dependent mitosis.

The success of PARP inhibitors in the treatment of homologous recombination–deficient tumors demonstrates the value of targeting cancer-specific liabilities ^78^. Given the widespread prevalence of chromosome imbalance in human cancers ^79^, mitotic vulnerabilities that arise selectively in chromosomally unstable cells may offer a similarly promising route for precision therapy. These cancers are characterized not only by altered chromosome number, but also by rewired mitotic control, including changes in SAC and APC/C activity. Our data suggest that loss of the SKA complex exposes this altered control as a cancer-specific vulnerability. Thus, we identify the SKA complex as a candidate therapeutic target in chromosomally unstable cancers. Chemical tools such as PROTACs or molecular glues may enable the targeting of this structural complex ^80^. As our analyses position the SKAc in a mitotic pathway distinct from KIF18A, SKAc inhibition may prove to be effective as a therapeutic option even in circumstances of resistance to KIF18A inhibition.

## Materials and Methods

### Cell culture

All RPE1 cells were grown in DMEM (Gibco) supplemented with 7.5% fetal bovine serum (FBS, Capricorn). HeLa cells were grown in Advanced DMEM/F-12 (Gibco) with 9% FBS and 1% GlutaMAX (Gibco). OVCAR-3 cells were grown in RPMI 1640 (Gibco) with 20% FBS and 10.6 μg/mL human insulin (Sigma-Aldrich). HCT-116 cells were grown in McCoy’s 5A (Modified) Medium (Gibco) with 9% FBS and 1% GlutaMAX (Gibco). DLD-1 and HCC-1806 cells were grown in RPMI 1640 (Gibco) with 9% FBS. All cell culture media were supplemented with 1% penicillin-streptomycin (Gibco). All cell lines were grown at 37°C under 5% CO_2_ and routinely tested negative for mycoplasma contamination. RPE1-hTERT-iCas9-Δ*TP53* cells were described previously ^81^. CRISPR-Cas9 genome editing was used to generate RPE1-hTERT-iCas9-Δ*TRIP13*Δ*TP53*, RPE1-hTERT-iCas9-Δ*SKA1/2/3*Δ*TRIP13*Δ*TP53* and RPE1-hTERT-iCas9-Δ*TP53*--Δ*KIF18A*Δ*TRIP13* cell lines. Edited cell pools were seeded at low density, after which single-cell clones were isolated and validated by TIDE analysis ^82^ and western blotting. Primer sequences are listed in Supplementary Table 1. HCT-116 cells were kindly provided by Geert Kops (Hubrecht Institute), and OVCAR-3 cells by Marcel van Vugt (University Medical Center Groningen).

### CRISPR-Cas9 genome editing

All genome editing was performed in cell lines containing an inducible Tet-On Cas9 expression system. One day before transfection, cells were seeded and Cas9 expression was induced with 200 ng/mL doxycycline. Transfections were performed using Lipofectamine RNAiMAX (Invitrogen) at a 1:1000 dilution with equimolar concentrations of crRNA and tracrRNA (IDT) in OptiMEM (Gibco), to a final concentration of 50 nM crRNA:tracrRNA duplex per 40,000 cells. crNRA sequences are listed in Supplementary Table 2.

### Lentiviral transduction

To generate the RPE1-hTERT-iCas9-Δ*TRIP13*Δ*TP53 +* pCMVie-TRIP13 cell line (referred to as Δ*TRIP13* + *pCMV-TRIP13*), TRIP13 was re-expressed via lentiviral transduction. TRIP13 cDNA was cloned into pLenti CMVie-IRES-BlastR, a gift from Ghassan Mouneimne (Addgene plasmid # 119863). Lentiviral particles were produced in HEK293T cells and used to transduce the RPE1-hTERT-iCas9-Δ*TP53*-Δ*TRIP13* cells. Transduced cells were selected with 10 μg/mL blasticidin and TRIP13 expression was confirmed by western blotting.

### RNA interference

siRNA-mediated knockdown of target genes was achieved by transfecting cells with a final concentration of 20 nM ON-TARGETplus siRNA SMARTPools (Dharmacon Reagents) using Lipofectamine RNAiMAX at a 1:1000 dilution. Analysis or re-seeding of cells was performed 48 hours post-transfection unless stated otherwise. Gene knockdown was validated by western blotting. Target sequences included *ANAPC15* (L-020237-02-0005), *CSNK1A1* (L-003957-00-0005), *CUL3* (L-010224-00-0005), *HAUS3* (L-018131-01-0005), *MAGOH* (L-011327-00-0005), *PLK1* (L-003290-00-0010), *POLR2L* (L-013133-01-0020), *PPP1CB* (L-008685-00-0005), *PPP1R12A* (L-011340-00), and *SKA2* (L-018781-01-0005). A non-targeting (NT) siRNA pool was used as a negative control (D-001810-01-20).

### Genome-wide CRISPR-Cas9 screen

The Δ*TRIP13* genome-wide CRISPR-Cas9 screen was performed in RPE1-hTERT-iCas9-Δ*TRIP13*Δ*TP53* cells, which form an isogenic pair to the RPE1-hTERT-iCas9-Δ*TP53* cells previously screened, as described in ^81^. The Δ*SKA1*Δ*TRIP13* screen was performed using RPE1-hTERT-iCas9-Δ*SKA1*Δ*TRIP13*Δ*TP53* and RPE1-hTERT-iCas9-Δ*TRIP13*Δ*TP53* cells. During the screens, all cells were grown in DMEM medium (Gibco) supplemented with 9% fetal bovine serum, 1% penicillin-streptomycin (Gibco), 1% sodium pyruvate (Gibco) and 0.5 μg/mL Amphotericin B (Gibco). The screens were performed using the TKOv3 library ^83^ at a 400-fold library representation, in triplicate. Cells were transduced with pLCKO-TKOv3 in medium containing 8 μg/mL polybrene at an MOI of approximately 0.2. Medium was refreshed after 24 hours, and 5 μg/mL puromycin was added to select for transduced cells. After three days of selection, cells were divided into three independent populations, harvested for the t=0 sample, and then reseeded in medium containing 200 ng/mL doxycycline to induce Cas9 expression. Cells were passaged every three population doublings, which corresponded to every three days, for a total of twelve population doublings. Doxycycline was maintained in the medium throughout culturing. Genomic DNA was isolated from cell pellets using the Blood and Cell Culture DNA Maxi Kit (Qiagen), and integrated sgRNA sequences were amplified by PCR using KAPA HiFi HotStart ReadyMix (Roche). Lentiviral inserts were amplified by PCR and Illumina adapters and i5 and i7 index sequences were added in two PCR steps as described previously ^81^. The triplicates from the different samples (t=0 and t=4) were pooled and sequenced on a NovaSeq6000 (Illumina). Sequences were mapped to the TKOv3 library (not allowing any mismatches) and the differential gRNA count data were analyzed and attributed to gene scores using IsogenicZ, an adapted version of DrugZ optimized for isogenic cell line screens, using a previous screen in the RPE1-Δ*TP53* parental cell line as a control ^81,84,85^.

### GOterm analysis

The hits identified in the CRISPR-Cas9 screen were analyzed for gene ontology (GO) enrichment using the GO Ontology database (DOI: 10.5281/zenodo.15066566, released 2025-03-16) ^86,87^ and https://string-db.org/ ^88,89^.

### Chemical compounds

All chemical compounds were dissolved in DMSO and used at the indicated concentrations unless otherwise specified. Nocodazole (Sigma-Aldrich M1404) was used at a final concentration of 0.3 μM for 18 hours to arrest cells in mitosis. Palbociclib (Selleck Chem S1116) was used at a final concentration of 200 nM for 24 hours to synchronize cells in G1 phase. RO-3306 (Sigma-Aldrich #217699) was used at 9 μM for 18 hours to synchronize cells at the G2/M transition. MG-132 (Sigma-Aldrich C2211) was used at a final concentration of 5 μM to arrest cells in anaphase. ProTAME (Sigma-Aldrich SML3977) was used in varying concentrations to inhibit the APC/C. Noscapine (Sigma-Aldrich 363960) was used in varying concentrations to disrupt microtubule assembly. Sovilnesib, a KIF18A inhibitor developed by Amgen and kindly provided to us by M. Luijsterburg (LUMC), was used at 125 nM to inhibit KIF18A activity unless stated otherwise. Reversine (Sigma-Aldrich R3904) was used in varying concentrations to inhibit MPS1 and thus the spindle assembly checkpoint.

### Viability and clonogenic assays

Cells were seeded two days after transfection with crRNA: tracrRNA duplexes. For viability assays, cells were seeded in 96-well plates at a density of 500 cells per well, with an exception for OVCAR-3 cells which were seeded at a density of 1,500 cells per well. Seven days post-transfection, the culture medium was replaced with medium containing 40 μM Resazurin (Sigma-Aldrich). After a four-hour incubation at 37°C, fluorescence was measured at 545 nm excitation/600 nm emission using a CLARIOstar Plus plate reader (BMG Labtech). Fluorescence measurements were corrected for background fluorescence from wells containing medium only. For clonogenic assays, cells were seeded two days post-transfection in 6-well plates at a density of 1000 cells per well. Twelve days after transfection, cells were stained using 0.5 mg/mL crystal violet solution (Merck) with 20% MeOH.

### Drug response assays

Cells were seeded in 96-well plates at a density of 500 cells per well, with an exception for OVCAR-3 cells which were seeded at a density of 1,500 cells per well, one day prior to drug treatment. Three days post treatment, the culture medium was refreshed, and fresh drug was added. Seven days after initial drug addition, the culture medium was replaced with medium containing 40 μM Resazurin (Sigma-Aldrich). After a four-hour incubation at 37°C, fluorescence was measured at 545 nm excitation/600 nm emission using a CLARIOstar Plus plate reader (BMG Labtech). Fluorescence measurements were corrected for background fluorescence from wells containing medium only.

### Western blotting

For western blotting, cells were lysed in lysis buffer (50 mM Tris-HCl pH 7.4, 150 mM NaCl, 1% Triton X-100 supplemented with fresh protease and phosphatase inhibitors) for ten minutes on ice and centrifuged at 4°C and 14,000 RPM for ten minutes. Proteins were separated on 4-15% Mini-PROTEAN TGX Precast Protein gels (Bio-Rad), followed by a semi-dry transfer to a nitrocellulose membrane (Bio-Rad). After blocking in 5% milk in TBS-T for 30 minutes, membranes were incubated at 4°C overnight with the following primary antibodies at the indicated dilutions in 5% milk in TBS-T: mouse α-HSP90_α/β_ (Biolegend 675402, 1:9000), rabbit α-KIF18A (Bethyl A301-080A, 1:1000), mouse α-MAD2 (Sigma-Aldrich MABE866, 1:1000), mouse α-P31COMET (Sigma-Aldrich MABE451, 1:2000), rabbit α-REV7 (Abcam Ab180579, 1:1000), rabbit α-TRIP13 (Bethyl A303-605A, 1:1000). Polyclonal rabbit α-SKA1, α-SKA2, and α-SKA3 antibodies (described in ^45^, 1:2000) were kindly gifted by Iain Cheeseman (Whitehead Institute for Biomedical Research). Following primary antibody incubation, membranes were incubated with HRP-conjugated secondary antibodies for one hour at room temperature: goat α-mouse (Cell Signaling 7074, 1:5000) and goat α-rabbit (Cell Signaling 7076, 1:5000). Proteins were visualized using Clarity Western ECL Substrate (Bio-Rad). Quantification of intensity levels was performed using ImageJ, taking the mean grey intensity for the relevant bands and subtracting background intensity.

### Immunofluorescence and image analysis

Cells were seeded on 35-mm glass-bottom dishes with #1.5 (0.17 mm) cover glass thickness (MatTek Corporation)in 1 mL DMEM supplemented as described above to reach 80% confluence the following day. Where noted, cells were treated for 1 hour prior to fixation with either DMSO or 250 nM Sovilnesib (MedChemExpress, HY-132840). Where indicated, cells were additionally treated 30 minutes prior to fixation with 5 μM MG-132 (Merck, M7449).

Cells stained for α-tubulin, CREST, and CENP-E were fixed for 2 min in ice-cold methanol (−20 °C). For staining of CREST, KIF18a, and Ska3, cells were first rinsed for 10 seconds with pre-warmed (37 °C) cell extraction buffer (100 mM PIPES, 1 mM MgCl₂, 1 mM EGTA, 0.5% Triton X-100), followed by methanol fixation as described above. After fixation and three 5-minute PBS washes, cells were permeabilized with 0.5% Triton X-100 in water for 30 min at room temperature. To block nonspecific binding, cells were incubated in 1 mL blocking buffer (2% normal goat serum, NGS; Thermo Fisher, 10000C) for 2 hours at room temperature. Cells were then washed three times for 5 min in PBS and incubated with 300 µL primary antibody solution overnight at 4 °C. Primary antibodies were diluted in PBS containing 0.1% Triton X-100 and 1% NGS. The following primary antibodies were used: rabbit anti-KIF18A (A301-080A, Bethyl Laboratories, 1:500), rabbit anti-CENP-E (C7488, Sigma, 1:500), mouse anti-SKA3 (sc-390326, Santa Cruz, 1:100), rat anti-α-tubulin YL1/2 (MA1-80017, Invitrogen, 1:500), and human anti-centromere (CREST) serum (15-234, Antibodies Incorporated, 1:500). After primary antibody incubation and a PBS wash, cells were incubated for 1 hour at room temperature with 500 µL secondary antibody solution diluted in PBS containing 0.1% Triton X-100 and 1% NGS. Secondary antibodies were selected according to the species of the primary antibodies. For CENP-E, CREST, and α-tubulin staining, the following secondary antibodies were used: donkey anti-rat Alexa Fluor 647 (Abcam, ab150155, 1:500), goat anti-human DyLight 594 (Abcam, ab96909, 1:1000), and goat anti-rabbit Alexa Fluor 488 (Abcam, ab150073, 1:500), respectively. For KIF18a, Ska3, and CREST staining, the following secondary antibodies were used: donkey anti-rabbit Alexa Fluor 594 (Abcam, ab150064, 1:1000), donkey anti-mouse Alexa Fluor 488 (Abcam, ab150105, 1:500), and goat anti-human Alexa Fluor 647 (Jackson ImmunoResearch, 109-605-003, 1:1000), respectively. This was followed by washing three times 10 minutes in PBS. Cells were imaged immediately after staining or stored at 4 °C protected from light for up to one week prior to imaging.

Cells were imaged using a Zeiss LSM 800 laser scanning confocal microscope equipped with a Plan-Apochromat 63×/1.40 NA oil immersion objective (DIC M27). Images were acquired using ZEN Blue software (Zeiss). For tubulin–CENP-E staining experiments, images were acquired at 512 × 512 pixel resolution with a lateral sampling of 0.0495 µm per pixel and a z-step size of 0.5 µm, generating 31 optical sections per stack. Fluorescence signals were recorded sequentially using Alexa Fluor 488 (tubulin), Alexa Fluor 594 (CENP-E), and Alexa Fluor 647 (CREST) channels. Imaging was performed using unidirectional scanning (scan speed 14) with line averaging set to 1, and images were acquired as 8-bit datasets. For KIF18a–Ska3 staining experiments, images were acquired at 488 × 488 pixel resolution with a lateral sampling of 0.041 µm per pixel and a z-step size of 0.5 µm, generating 31 optical sections per stack. Fluorescence signals were recorded sequentially using Alexa Fluor 488 (Ska3), Alexa Fluor 594 (KIF18a), and Alexa Fluor 647 (CREST) channels. Imaging parameters were otherwise identical to those described above, and images were acquired as 16-bit datasets. Image stacks from these experiments were subsequently processed using the ZEN LSM Plus deconvolution algorithm (AutoWiener filter) prior to analysis. Z-stacks spanning approximately 15 µm were collected sequentially at multiple stage positions.

Image analysis was performed in Fiji. Spindle length was measured in maximum-intensity projection images as the distance between the tubulin foci corresponding to the two spindle poles. Multipolar and highly tilted spindles were not imaged. In the analyzed dataset, spindle poles were separated by no more than eight z-planes. Maximum metaphase plate spread was quantified as the distance between the outermost CREST signals along the main spindle axis. A fixed-size oval region of interest (ROI) was used for all measurements, with the ROI size corresponding to the CREST kinetochore signal. Background intensity was measured in the cytoplasm at three independent locations, averaged, and subtracted from the kinetochore signal. For full width at half maximum (FWHM) measurements, a Gaussian blur (σ = 1) was applied before analysis. To quantify kinetochore alignment along the spindle axis using an independent metric, intensity profiles of CREST signals were analyzed in Fiji, as described before ^90^. Briefly, a 200-pixel linear ROI was drawn along the main spindle axis, and gray-value intensities were extracted. Intensity profiles were centered on the metaphase plate. Spatial coordinates were converted from pixels to micrometers (µm) prior to analysis. The FWHM of the CREST signal was used as a quantitative measure of kinetochore alignment. For each profile, the half-maximum threshold was defined as yₕₘ = (yₘₐₓ + yₘᵢₙ)/2, where yₘₐₓ represents the global maximum intensity and yₘᵢₙ the local background baseline. The positions at which the signal crossed this threshold (xₗₑ_f_ₜ and xᵣᵢ_g_ₕₜ) were determined by linear interpolation between adjacent data points to obtain sub-pixel accuracy. The FWHM was then calculated as FWHM = xᵣᵢ_g_ₕₜ − xₗₑ_f_ₜ. Smaller FWHM values indicate tighter kinetochore clustering at the metaphase plate, whereas larger FWHM values or multimodal profiles indicate chromosome misalignment or spindle disorganization.

### Flow cytometry

Upon harvesting the cells and culture supernatant, cells were rinsed in cold PBS with 0.05% EDTA, trypsinized, and washed with PBS before fixation in cold 70% ethanol. After incubation at -20°C for at least an hour, cells were permeabilized with 0.05% Tween-20 in PBS. Following a wash with 0.05% Tween-20 in PBS, cells were blocked and stained overnight at 4°C using α-Histone H3 phospho-Serine 10-Alexa Fluor 647 (Biolegend 650806) diluted 1:100 in 0.05% Tween-20/2% BSA in PBS. After three washes with 0.05% Tween-20 in PBS, the cells were stained with 0.5 μg/mL DAPI in 0.05% Tween-20/2% BSA in PBS. Cells were analyzed using an LSRFortessa X-20 (BD Biosciences) or CytoFLEX LX Flow Cytometer (Beckman Coulter). FlowJo software was used for data processing. To assess the fraction of mitotic cells, live-cells were first gated, followed by single cells gating. These were then gated for the pH3-positive, 4N population.

### Live-cell imaging

Cells were grown in 8-well Nunc Lab-Tek II Chambered Coverglasses (Thermo Scientific), with the addition of the appropriate chemical compounds needed for the experiment. Four hours prior to the start of imaging, SPY650-DNA (Spirochrome, 1:2000) and verapamil (Spirochrome, 1:10,000) were added to the cells. Imaging was performed every four minutes using a THUNDER Imager Live-cell (Leica Microsystems) equipped with a HC PL APO 20x/0.80 objective (dry) and a deep-cooled 4.2 MP sCMOS camera, with cells maintained at 37°C under 5% CO_2_. Image analysis was performed in Fiji based on maximum-intensity projection images. All scoring was performed blinded.

### Chromosome spreads

Cells were treated with 200 ng/mL demecolcin for 20 minutes, harvested, and resuspended in 0.075 M KCl for 20 minutes. Subsequently, cells were fixed in methanol:acetic acid (3:1) and dropped onto glass slides. Following staining with 5% Giemsa solution, metaphases were assessed for chromosome number per cell.

### Single cell karyotyping

Sequencing was performed using a NextSeq 2000 (Illumina; up to 74 cycles), after which the generated data were demultiplexed using sample-specific barcodes and changed into fastq files using bcl2fastq (Illumina; version 1.8.4). Reads were aligned to the human reference genome (GRCh38/hg38) using Bowtie2 (version 2.2.4; ^91^) and duplicate reads were marked with BamUtil (version 1.0.3; ^92^). Copy number variation analysis was performed with AneuFinder (version 1.30.0; https://github.com/ataudt/aneufinder; ^93^). After GC correction and blacklisting of artefact-prone regions (extreme low or high coverage in control samples), libraries were analyzed using the dnacopy and edivisive copy number calling algorithms with variable width bins (average bin size = 1 Mb; step size = 500 kb) and breakpoint refinement (confint = 0.95). Blacklists and variable width bins were constructed using an euploid reference ^94^. Results were curated by requiring a minimum concordance of 90% between the results of the two algorithms. Libraries with on average less than 10 reads per chromosome copy of each bin (2-somy: 20 reads, 3-somy: 30 reads, etc.) were discarded. This minimum number of reads is approximately 55,000 for a diploid genome. The aneuploidy score of each bin was calculated as the absolute difference between the observed copy number and the euploid copy number. The weighted average of the bins is the score per library, which was averaged to the score per sample. The heterogeneity score of each bin was calculated as the proportion of pairwise comparisons (cell 1 versus cell 2, cell 1 versus cell 3, etc.) that showed a difference in copy number (*e.g*., cell 1: 2-somy and cell 2: 3-somy). The score of each sample is the weighted average of all bin scores.

### DepMAP analyses

For gene essentiality assessment, gene effect scores from the DepMap project were downloaded and analyzed (^95^ and (https://depmap.org/portal/)). Gene effect scores quantify how essential a gene is for a cancer cell line. A score close to -1 indicates a gene is likely essential, which means knocking it out significantly reduces cell viability. A score near 0 or above indicates that the gene is likely non-essential. Gene effects scores for Δ*TRIP13* SV genes were compared with the gene effect scores of all screened genes and with those that are classified as ‘common essential’ in DepMap.

Aneuploidy scores (AS) for human cancer cell lines were obtained from ^96^. mRNA gene expression values and CRISPR dependency scores (DS; Chronos) were obtained from the DepMap25Q2release ^38^ (OmicsExpressionExpectedCountHumanAllGenes.csv, and CRISPRGeneEffect.csv, respectively). FPKM values were calculated by normalizing the read counts for each gene to both the total library size (in millions of mapped reads) and the gene length (in kilobases), as defined in the Ensembl Homo Sapiens reference annotation (GRCh38, release 112), following standard protocols. In analyses where cells were divided into aneuploid and near-euploid groups, the division was performed using the top and bottom quartiles of AS (Top: AS ≥ 21; Bottom: AS ≤ 8). In analyses where cells were divided into KIF18A-dependent and -resistant groups, the division was performed using the top and bottom quartiles of KIF18A DS (Top: DS ≤ –1.73567; Bottom: DS ≥ –0.76964). Gene preferential essentiality in aneuploid versus near-euploid cells was calculated as the mean DS difference between groups, and associated P-values were derived from unpaired, two-tailed t-tests. Correlations between gene dependency and aneuploidy were assessed using the Spearman’s rank correlation coefficient (ρ), with P-values derived from two-tailed tests. Co-dependency analysis was performed using pairwise Spearman’s correlation coefficients (ρ); corresponding P-values were computed using two-tailed tests and adjusted for multiple testing using the Benjamini–Hochberg false discovery rate (FDR) procedure to obtain Q-values. Results are displayed as –log₁₀(Q-value) versus the correlation coefficient itself. Gene expression association with KIF18A dependency was evaluated by comparing mean mRNA expression (FPKM) between dependent and resistant groups, with P-values derived from unpaired, two-tailed t-tests. Effect sizes were quantified using Cohen’s d, calculated as the difference in means divided by the pooled standard deviation. In analyses assessing the *SKA1*, *SKA2*, and *SKA3* genes as a complex, SKA-complex expression was calculated as the average FPKM mRNA expression of the three genes in each cell line.

## Acknowledgments

We thank Isabel Potters (Princess Máxima Center for pediatric oncology) for computational support, Iain Cheeseman (Whitehead Institute for Biomedical Research, Cambridge, USA), Geert Kops (Hubrecht Institute, Utrecht, The Netherlands), Marcel van Vugt (UMC Groningen, Groningen, The Netherlands) and Andrea Musacchio (Max Planck Institute of molecular physiology, Dortmund, Germany) for reagents, and Marialucrezia Losito (IPSomics) for scWGS analysis. We acknowledge Stefano Maffini (Max Planck Institute of molecular physiology) and Marco Novais-Cruz (Princess Máxima Center for pediatric oncology) for discussions. We thank members of the Medema-Vader lab for input and support, and Pim Huis in ‘t Veld (Max Perutz Labs, Vienna, Austria) for discussions and *in silico* protein complex modeling. We thank Martijn Luijsterburg and Sjoerd Klaasen (LUMC, Leiden, The Netherlands) for sharing unpublished observations and for Sovilnesib. We acknowledge Marcel van Vugt, Andreas Hochwagen (NYU, New York, USA) and Valentina Piano (University of Cologne, Cologne, Germany) for input on the manuscript.

## Funding

Amsterdam UMC AUMC research fellowship (GV)

Amsterdam UMC ADORE2024-9-01 grant (RMF)

Croatian Science Foundation Grant IP-2024-05-5336 (IMT)

Croatian Science Foundation Grant IPCH-2022-10-9344 (IMT)

Dutch Cancer Society KWF exploration grant 15228 (GV)

European Research Council Starting Grant 945674 (UBD)

European Research Council Synergy Grant 855158 (IMT)

Israel Cancer Association Grant 20230018 (UBD)

Oncode Institute (RHM)

Scientific Center of Excellence - QuantiXLie Grant PK.1.1.10.0004 (IMT)

## Author contributions

Experimentation: MSU, ESC, KV, DR, KdL, MAR, and TDK

Conceptualization: MSU, GV, UBD, and RHM

Data analysis and computation: MSU, KdL, AT, IJH

Supervision: GV, UBD, RMFW, RDL, IMT, and FF

Project management: MSU, GV, UBD, RMFW, RDL

Funding acquisition: GV, RHM, IMT, UBD, RMFW, and FF

Writing: MSU and GV with input from all authors

## Data, code, and materials availability

All data are available in the main text or supplementary materials. Materials are available from the corresponding author as applicable.

**Movie S1. RPE1 *ΔTP53* (WT) unperturbed: completed mitosis**

Representative time-lapse video of an unperturbed RPE1-*hTERT*-iCas9 Δ*TP53* (WT) cell, stained with a DNA dye. The cell goes through and completes mitosis. Time is indicated as hours:minutes, scale bar = 10 μm.

**Movie S2. RPE1 *ΔTRIP13ΔTP53* unperturbed: completed mitosis**

Representative time-lapse video of an unperturbed RPE1-*hTERT*-iCas9 Δ*TRIP13*Δ*TP53* cell, stained with a DNA dye. A. The cell goes through and completes mitosis. Time is indicated in hours:minutes, scale bar = 10 μm.

**Movie S3. RPE1 ΔTP53 (WT) siMock: completed mitosis**

Representative time-lapse video of a RPE1-*hTERT*-iCas9 Δ*TP53* (WT) cell, stained with a DNA dye, treated with mock siRNA. The cell goes through and completes mitosis. Time is indicated as hours:minutes, scale bar = 10 μm.

**Movie S4. RPE1 ΔTP53 (WT) siSKA2: completed mitosis**

Representative time-lapse video of a RPE1-*hTERT*-iCas9 Δ*TP53* (WT) cell, stained with a DNA dye, treated with *SKA2* siRNA. The cell goes through and completes mitosis. Time is indicated as hours:minutes, scale bar = 10 μm.

**Movie S5. RPE1 ΔTP53 (WT) siSKA2: aberrant anaphase**

Representative time-lapse video of a RPE1-*hTERT*-iCas9 Δ*TP53* (WT) cell, stained with a DNA dye, treated with *SKA2* siRNA. The cell goes through mitosis, which results in an aberrant anaphase. Time is indicated in hours:minutes, scale bar = 10 μm.

**Movie S6. RPE1 *ΔTP53* (WT) si*SKA2*: no mitotic exit**

Representative time-lapse video of a RPE1-*hTERT*-iCas9 Δ*TP53* (WT) cell, stained with a DNA dye, treated with *SKA2* siRNA. The cell enters mitosis and does not exit mitosis within the span of the video. Time is indicated in hours:minutes, scale bar = 10 μm.

**Movie S7. RPE1 *ΔTRIP13ΔTP53* siMock: completed mitosis**

Representative time-lapse video of a RPE1-*hTERT*-iCas9 Δ*TRIP13*Δ*TP533* cell, stained with a DNA dye, treated with mock siRNA. The cell goes through and completes mitosis. Time is indicated as hours:minutes, scale bar = 10 μm.

**Movie S8. RPE1 *ΔTRIP13ΔTP53* siMock: aberrant anaphase**

Representative time-lapse video of a RPE1-*hTERT*-iCas9 Δ*TRIP13*Δ*TP53*cell, stained with a DNA dye, treated with mock siRNA. The cell goes through mitosis, which results in an aberrant anaphase. Time is indicated as hours:minutes, scale bar = 10 μm.

**Movie S9. RPE1 *ΔTRIP13ΔTP53* si*SKA2*: completed mitosis**

Representative time-lapse video of a RPE1-*hTERT*-iCas9 Δ*TRIP13*Δ*TP53* cell, stained with a DNA dye, treated with *SKA2* siRNA. The cell goes through and completes mitosis. Time is indicated as hours:minutes, scale bar = 10 μm.

**Movie S10. RPE1 *ΔTRIP13ΔTP53* si*SKA2*: aberrant anaphase**

Representative time-lapse video of a RPE1-*hTERT*-iCas9 Δ*TRIP13*Δ*TP53* cell, stained with a DNA dye, treated with *SKA2* siRNA. The cell goes through mitosis, which results in an aberrant anaphase. Time is indicated as hours:minutes, scale bar = 10 μm.

**Movie S11. RPE1 ΔTP53 (WT) unperturbed: completed mitosis**

Representative time-lapse video of an unperturbed RPE1-*hTERT*-iCas9 Δ*TP53* (WT) cell, stained with a DNA dye. The cell goes through and completes mitosis. Time is indicated as hours:minutes, scale bar = 10 μm.

**Movie S12. RPE1 *ΔTP53*Δ*TRIP13* unperturbed: completed mitosis**

Representative time-lapse video of an unperturbed RPE1-*hTERT*-iCas9 *ΔTRIP13*Δ*TP53*cell, stained with a DNA dye. The cell goes through and completes mitosis. Time is indicated as hours:minutes, scale bar = 10 μm.

**Movie S13. RPE1 *ΔSKA2ΔTRIP13ΔTP53* unperturbed: completed mitosis**

Representative time-lapse video of an unperturbed RPE1-*hTERT*-iCas9 Δ*SKA2ΔTRIP13* Δ*TP53*cell, stained with a DNA dye. The cell goes through and completes mitosis. Time is indicated as hours:minutes, scale bar = 10 μm.

**Movie S14. RPE1 *ΔSKA2ΔTRIP13ΔTP53* unperturbed: aberrant anaphase**

Representative time-lapse video of an unperturbed RPE1-*hTERT*-iCas9 Δ*SKA2ΔTRIP13*Δ*TP53* cell, stained with a DNA dye. The cell goes through mitosis, which results in an aberrant anaphase. Time is indicated as hours:minutes, scale bar = 10 μm.

**Movie S15. RPE1 *ΔSKA3ΔTRIP13ΔTP53*: completed mitosis**

Representative time-lapse video of an unperturbed RPE1-*hTERT*-iCas9 *ΔSKA3ΔTRIP13ΔTP53* cell, stained with a DNA dye. The cell goes through and completes mitosis. Time is indicated as hours:minutes, scale bar = 10 μm.

**Movie S16. RPE1 *ΔSKA3ΔTRIP13ΔTP53*: aberrant anaphase**

Representative time-lapse video of an unperturbed RPE1-*hTERT*-iCas9 *ΔSKA3ΔTRIP13ΔTP53* cell, stained with a DNA dye. The cell goes through mitosis, which results in an aberrant anaphase. Time is indicated as hours:minutes, scale bar = 10 μm.

**Movie S17. RPE1 *ΔSKA2ΔTRIP13ΔTP53* + ProTAME: aberrant anaphase**

Representative time-lapse video of an RPE1-*hTERT*-iCas9 Δ*SKA2ΔTRIP13*Δ*TP53*cell, stained with a DNA dye and treated with 2 μM ProTAME. The cell goes through mitosis, which results in an aberrant anaphase. Time is indicated as hours:minutes, scale bar = 10 μm.

**Movie S18. RPE1 *ΔSKA2ΔTRIP13ΔTP53* + ProTAME: aberrant anaphase**

Representative time-lapse video of an RPE1-*hTERT*-iCas9 *ΔSKA2ΔTRIP13ΔTP53* cell, stained with a DNA dye and treated with 2 μM ProTAME. The cell goes through mitosis, which results in an aberrant anaphase. Time is indicated as hours:minutes, scale bar = 10 μm.

**Data S1.**

(**A**) Raw counts from the genome-wide CRISPR-Cas9 screen in RPE1-*hTERT*-iCas9 *ΔTP53* and RPE1-*hTERT*-iCas9 *ΔTRIP13ΔTP53* cells. (**B**) Raw counts for statistically significant synthetic viable hits from the genome-wide CRISPPR-Cas9 screen in RPE1-*hTERT*-iCas9 *ΔTP53* and RPE1-*hTERT*-iCas9 *ΔTRIP13ΔTP53* cells. (**C**) Raw counts for statistically significant synthetic lethal hits from the genome-wide CRISPPR-Cas9 screen in RPE1-*hTERT*-iCas9 *ΔTP53* and RPE1-*hTERT*-iCas9 *ΔTRIP13ΔTP53* cells. (**D**) Raw counts for genes of interest from the genome-wide CRISPR-Cas9 screen in RPE1-*hTERT*-iCas9 *ΔTRIP13ΔTP53* and RPE1-*hTERT*-iCas9 *ΔSKA1ΔTRIP13ΔTP53* cells.

---

**Fig. S1.**
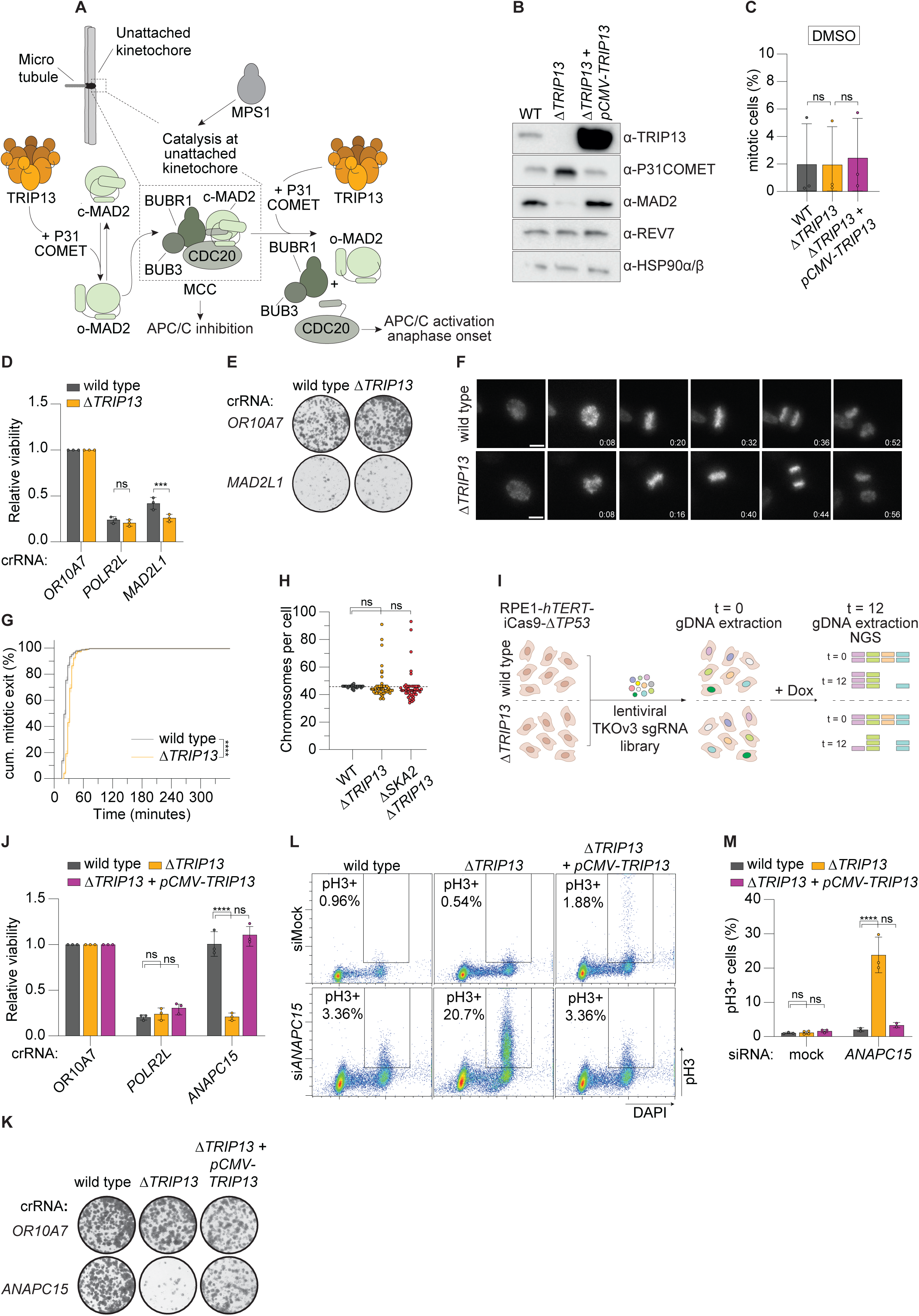
(**A**) Schematic depicting the mitotic checkpoint complex and the role of TRIP13 in regulation of this complex by converting c-MAD2 to o-MAD2. (**B**) Representative western blot of TRIP13, P31^COMET^, MAD2, and REV7 in RPE1-*hTERT*-iCas9 Δ*TP53* (WT), RPE1-*hTERT*-iCas9 Δ*TRIP13*Δ*TP53* (Δ*TRIP13*), and RPE1-*hTERT*-iCas9 Δ*TRIP13*Δ*TP53 + pCMV-TRIP13* (Δ*TRIP13 +* p*CMV-TRIP13*) cell lines. (**C**) Percentage of pH3-positive mitotic cells measured by flow cytometry following 18-hour DMSO treatment. (**D**) Relative viability upon CRISPR-based gene knockout. (**E**) Representative clonogenic survival assay upon CRISPR-based gene knockout. (**F**) Representative images of WT and Δ*TRIP13* cells stained with a live-cell DNA probe, undergoing division. Time is indicated in hours:minutes. Scale bar, 10 μm. (**G**) Mitotic exit timing of unperturbed WT and Δ*TRIP13* cells. Cells were tracked for more than 20 hours, but no new mitotic exits were observed after 6 hours. Only cells observed entering mitosis were included in this analysis. Shaded area indicates standard error. 229 WT and 221 Δ*TRIP13* cells were analyzed. (**H**) Chromosome counts of WT, Δ*TRIP13*, and RPE1-*hTERT*-iCas9 Δ*SKA2*Δ*TRIP13*Δ*TP53* (Δ*SKA2* Δ*TRIP13)* cells. *n* = 50 cells per condition. (**I**) Schematic overview of the genome-wide CRISPR-Cas9 knockout screen workflow performed in triplicate. Dox, doxycycline; NGS, next-generation sequencing. (**J**) Relative viability upon CRISPR-based gene knockout. (**K**) Representative clonogenic survival assay upon CRISPR-based gene knockout. (**L**) Representative flow cytometry plots showing gating used to identify mitotic populations (phospho-Histone H3-ser10-positive, 4N DNA content) after treatment with mock or *ANAPC15* siRNA. (**M**) Percentage of phospho-Histone H3-ser10-positive mitotic cells measured by flow cytometry after treatment with mock or *ANAPC15* siRNA. Data in (C-D), (J), and (M) were obtained from three biological independent experiments with technical duplicates (C) and (M) or triplicates (D) and (J). Statistical significance was assessed by one-way ANOVA (C) and (H), two-way ANOVA with correction for multiple comparisons (D), (J), and (M), and log-rank test (G). ***P < 0.001; ****P < 0.0001; ns, not significant.

**Fig. S2.**
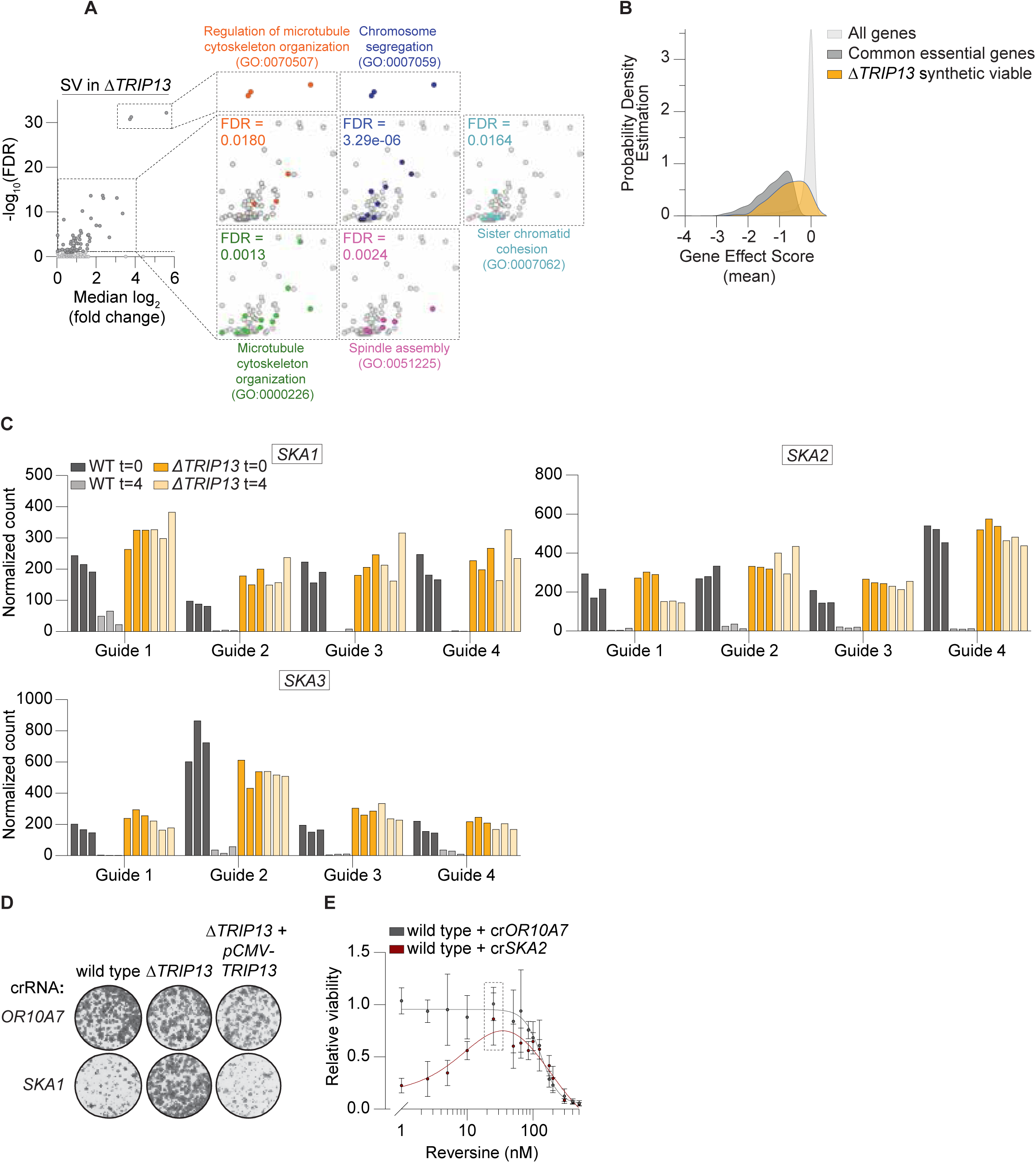
(**A**) Visual representation of synthetic viable (SV) genes involved in biological processes captured by five over-represented Gene Ontology (GO) terms within the total group of SV hits. Statistical significance of enriched GO terms is indicated by false discovery rate (FDR) values. (**B**) Probability density distribution of gene effect scores from DepMap for synthetic viable (SV) hits from our Δ*TRIP13* screen (orange), common essential genes (dark grey), and all genes (light grey). The probability density estimation reflects the relative frequency of gene effect scores. **(C)** Normalized counts of recovered *SKA1-3* sgRNAs at the first and last time point of the genome-wide CRISPR-Cas9 knockout screens in WT and Δ*TRIP13* cells. Each condition was assessed in triplicate. **(D)** Representative clonogenic survival assay of RPE1-*hTERT*-iCas9 Δ*TP53* (WT), RPE1-*hTERT*-iCas9 Δ*TRIP13*Δ*TP53* (Δ*TRIP13*), and RPE1-*hTERT*-iCas9 Δ*TRIP13*Δ*TP53 + pCMV-TRIP13* (Δ*TRIP13 +* p*CMV-TRIP13*) cell lines upon CRISPR-based gene knockout. **(E)** Relative viability after CRISPR-based gene knockout and treatment with different concentrations of Reversine. Results obtained for the concentration of 25 nM, indicated with the dashed lines, are also shown in Fig. 1E. Data were obtained from 3 biologically independent experiments with technical triplicates.

**Fig. S3.**
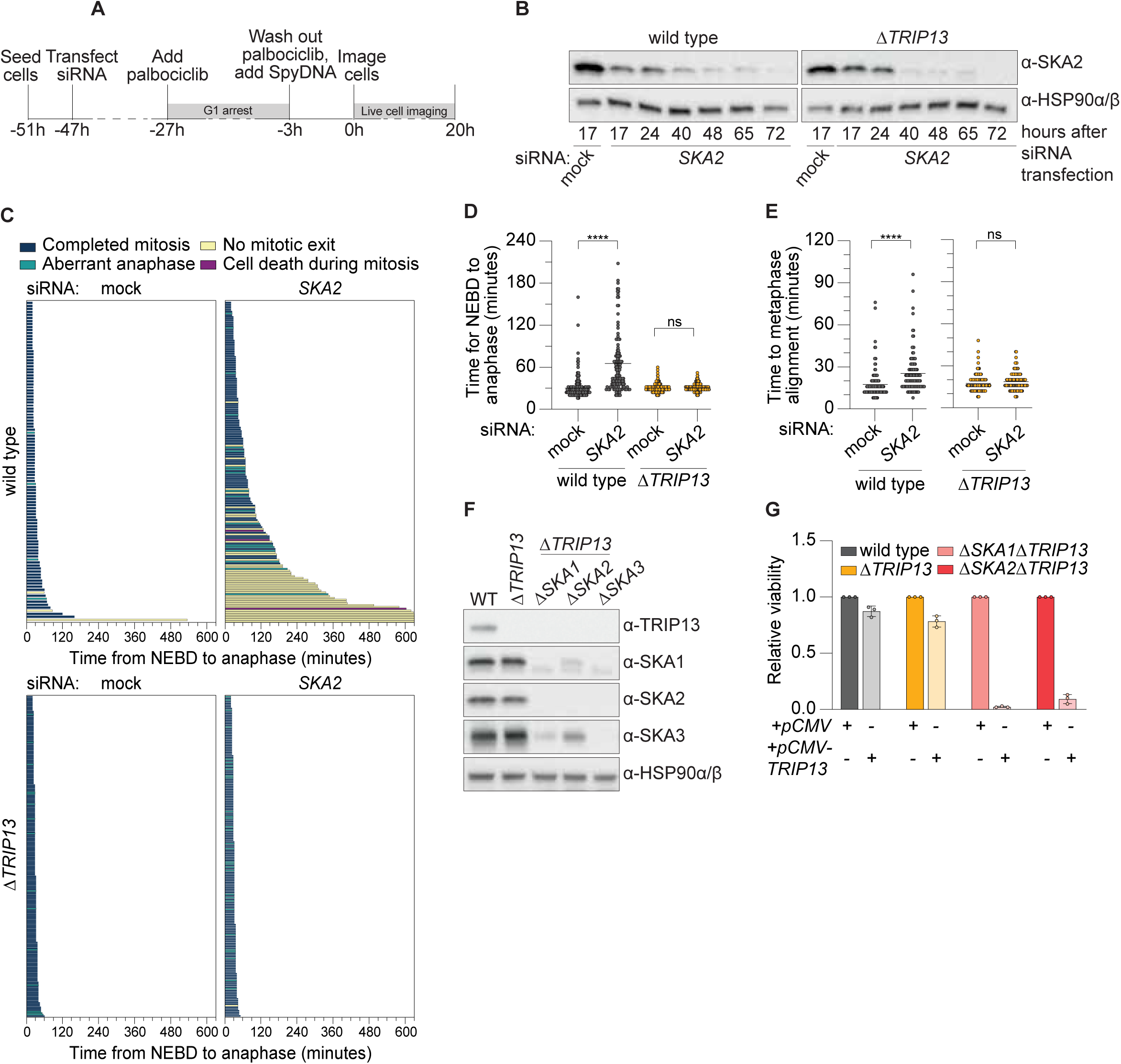
**(A)** Schematic overview of the live-cell imaging workflow used to image the first mitosis after protein degradation by siRNA. (**B**) Western blot of SKA2 in RPE1-*hTERT*-iCas9 Δ*TP53* (WT) and RPE1-*hTERT*-iCas9 Δ*TRIP13*Δ*TP53* (Δ*TRIP13*) cells, at different time points after treatment with mock or *SKA2* siRNA. (**C**) Single-cell mitotic duration (measured as time from nuclear envelope breakdown (NEBD) to anaphase) of WT and Δ*TRIP13* cells treated with mock or *SKA2* siRNA. Cell fates are color-coded: blue bars indicate completed mitosis, yellow bars indicate lack of mitotic exit during the duration of the video, teal bars indicate an aberrant anaphase, purple bars indicate mitotic cell death. Rows represent individual cells. The x-axis is truncated at 630 minutes, five WT si*SKA2* cells were filmed longer without exiting mitosis (764, 784, 804, 920, and 1060 minutes). For (C) and (D), 169, 178, 221, and 195 cells were analyzed for WT siMock, WT siSKA2, Δ*TRIP13* siMock, and Δ*TRIP13* siSKA2 cells, respectively. (**D**) Mitotic duration (measured as time from nuclear envelope breakdown (NEBD) to anaphase) of WT and Δ*TRIP13* cells treated with mock or *SKA2* siRNA. Only cells observed entering and exiting mitosis were included in this analysis. (**E**) Time to metaphase alignment of WT and Δ*TRIP13* cells treated with mock or *SKA2* siRNA. 134, 167, 179, and 163 cells were analyzed for WT siMock, WT siSKA2, Δ*TRIP13* siMock, and Δ*TRIP13* siSKA2 cells, respectively. (**F**) Representative western blot of TRIP13, SKA1, SKA2, and SKA3 in WT, Δ*TRIP13*, and RPE1-*hTERT*-iCas9 Δ*SKA1/2/3*Δ*TRIP13*Δ*TP53* (Δ*SKA1*Δ*TRIP13,* Δ*SKA2*Δ*TRIP13*, Δ*SKA3*Δ*TRIP13*) cells. (**G**) Relative viability after transfection with *pCMV*-*TRIP13* or an empty vector (*pCMV*). Data in (C), (D), and (G) were obtained from three biological independent experiments, for (G) with technical triplicates. Data in (E) were obtained from two or three biological independent experiments. Statistical significance was assessed by two-way ANOVA with correction for multiple comparisons for (D) and (E). ****P < 0.0001; ns, not significant.

**Fig. S4.**
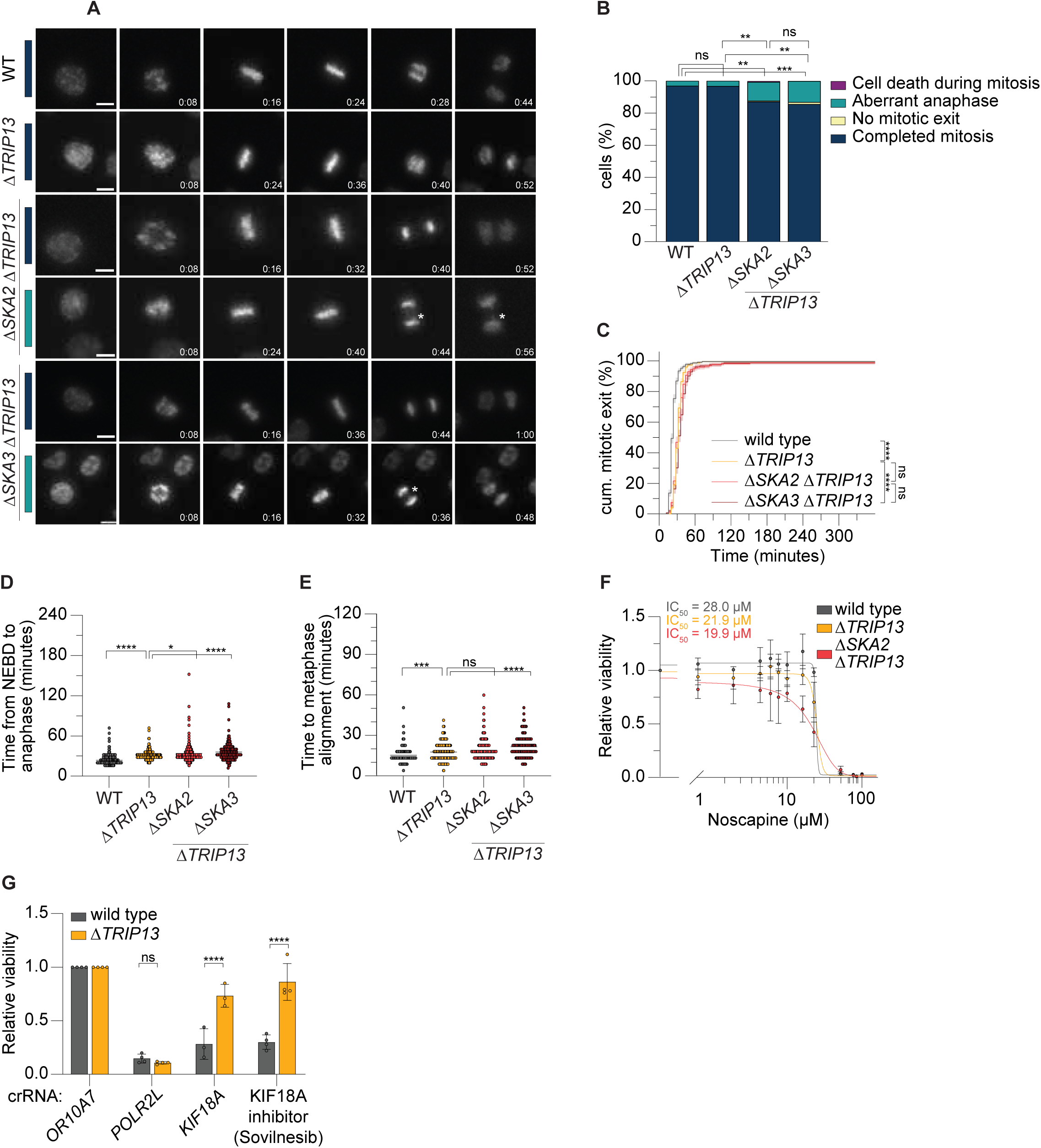
(**A**) Representative images of RPE1-*hTERT*-iCas9 Δ*TP53* (WT), RPE1-*hTERT*-iCas9 Δ*TRIP13*Δ*TP53* (Δ*TRIP13*), RPE1-*hTERT*-iCas9 Δ*SKA2*Δ*TRIP13*Δ*TP53* (Δ*SKA2*Δ*TRIP13*), and RPE1-*hTERT*-iCas9 Δ*SKA3*Δ*TRIP13*Δ*TP53* (Δ*SKA3*Δ*TRIP13*) cells stained with a live-cell DNA probe, undergoing division. Cell fates are color-coded: blue bars indicate completed mitosis, teal bars indicate an aberrant anaphase. An asterisk is placed to identify aberrancies. Time is indicated in hours:minutes. Scale bar, 10 μm. (**B**) Quantification of mitotic cell fates of WT, Δ*TRIP13*, Δ*SKA2*Δ*TRIP13*, and Δ*SKA3*Δ*TRIP13* cells by live-cell imaging. For (B-E), 229, 221, 171, and 155 were analyzed for WT, Δ*TRIP13*, Δ*SKA2*Δ*TRIP13*, and Δ*SKA3*Δ*TRIP13* cells, respectively. (**C**) Mitotic exit timing. Cells were tracked for more than 20 hours, but no new mitotic exits were observed after 6 hours. Only cells observed entering mitosis were included in this analysis. Shaded area indicates standard error. Data for WT and Δ*TRIP13* are identical to those shown in Figure S1G. (**D**) Mitotic duration (measured as time from nuclear envelope breakdown (NEBD) to anaphase). Only cells observed entering and exiting mitosis were included in this analysis. (**E**) Time to metaphase alignment of WT, Δ*TRIP13,* Δ*SKA2*Δ*TRIP13*, and Δ*SKA3*Δ*TRIP13* cells. (**F**) Dose-response relative viability curve after treatment with varying concentrations of Noscapine. (**G**) Relative viability upon CRISPR-based gene knockout or chemical inhibition of KIF18A using Sovilnesib. Data in (B-G) were obtained from three biological independent experiments, for (F) and (G) with technical triplicates. Statistical significance was assessed by Fisher’s exact test (B), log-rank test (C), or two-way ANOVA (D-E) and (G), each with correction for multiple comparisons. *P<0.05; **P < 0.01; ***P < 0.001; ****P < 0.0001; ns, not significant.

**Fig. S5.**
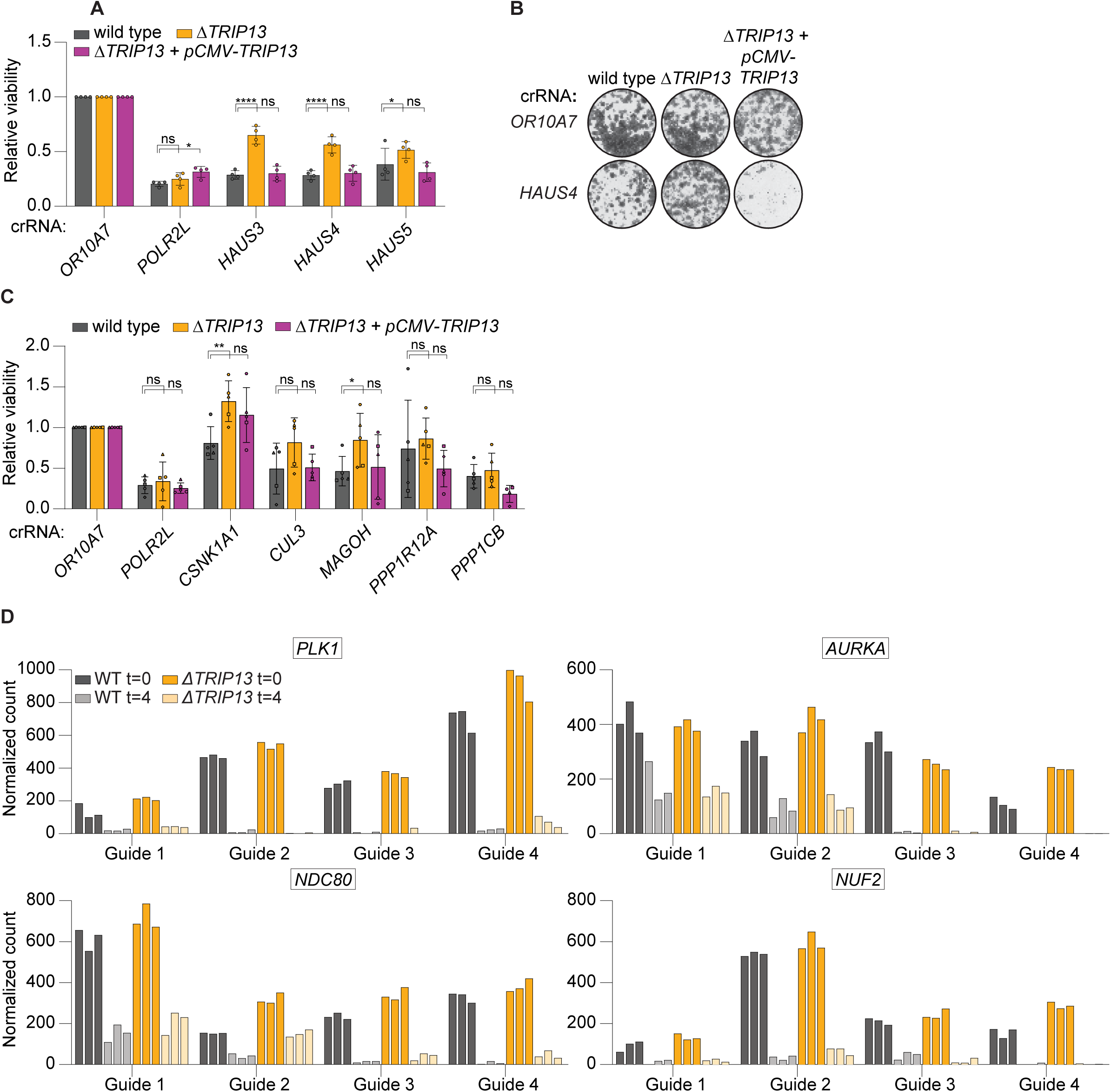
(**A**) Relative viability of RPE1-*hTERT*-iCas9 Δ*TP53* (WT), RPE1-*hTERT*-iCas9 Δ*TRIP13*Δ*TP53* (Δ*TRIP13*), and RPE1-*hTERT*-iCas9 Δ*TRIP13*Δ*TP53 + pCMV-TRIP13* (Δ*TRIP13 +* p*CMV-TRIP13*) cells upon CRISPR-based gene knockout. (**B**) Representative clonogenic survival assay upon CRISPR-based gene knockout. (**C**) Relative viability upon CRISPR-based gene knockout. (**D**) Normalized counts of recovered *PLK1*, *AURKA*, *NDC80*, and *NUF2* sgRNAs at the first and last time point of the genome-wide CRISPR-Cas9 knockout screens in WT and Δ*TRIP13* cells. Each condition was assessed in triplicate. Data in (A) and (C) were obtained from three biologically independent experiments with technical triplicates. Statistical significance was assessed by two-way ANOVA with correction for multiple comparisons. *P < 0.05; **P < 0.01; ****P < 0.0001; ns, not significant.

**Fig. S6.**
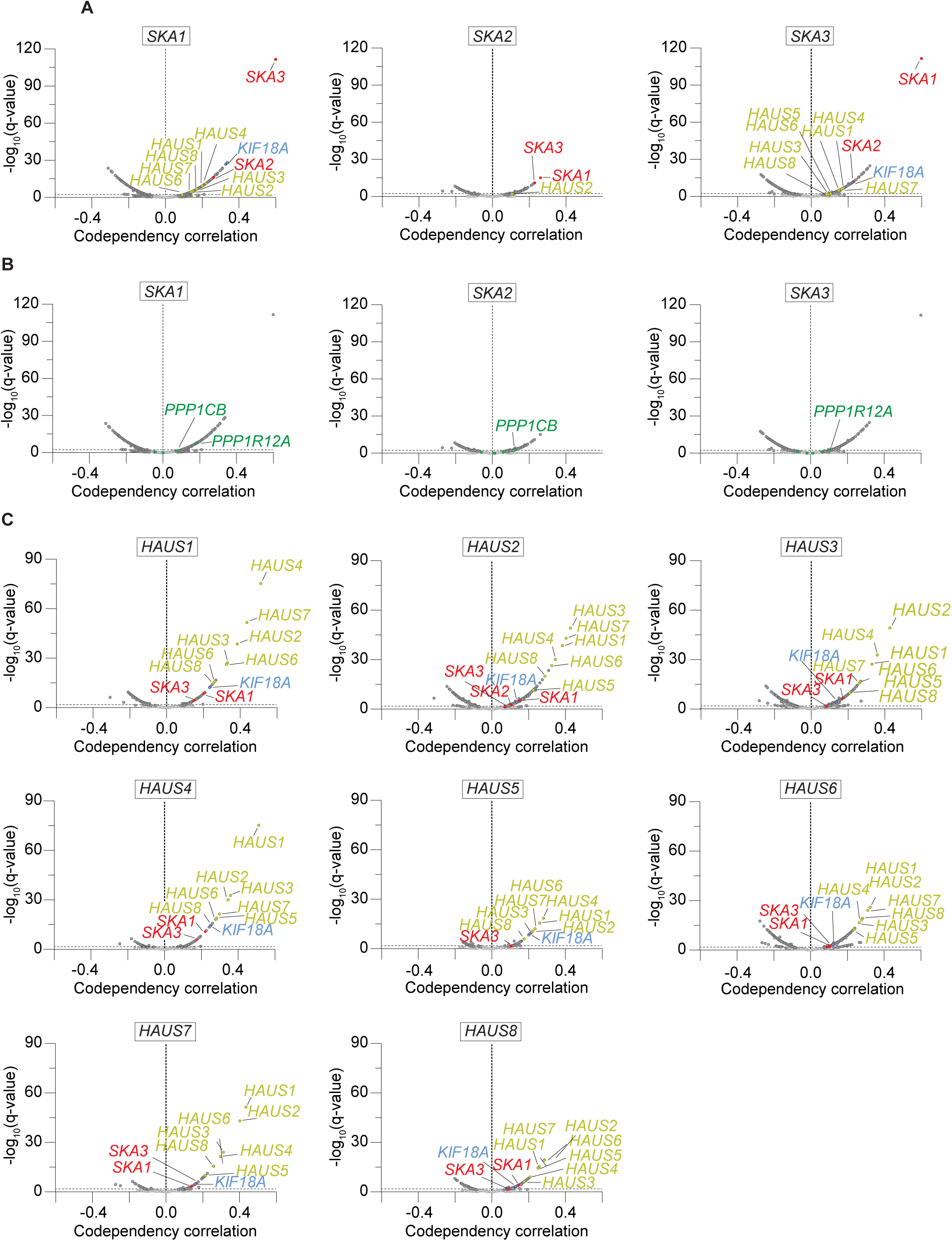
Volcano plots showing gene co-dependency correlations with *SKA1*, *SKA2* and *SKA3* across cancer cell lines, (**A**) specifically focused on the SKAc, KIF18A, and Augmin complex or (**B**) specifically focused on the other mitosis-linked synthetic viable hits identified in the CRISPR screen. The x-axis represents the Pearson correlation coefficient (r), the y-axis shows the significance. Genes with significant co-dependency (q < 0.05) are shown in dark gray. (**C**) Volcano plots showing gene co-dependency correlations with *HAUS1-8* across cancer cell lines, specifically focused on the SKAc, KIF18A, and the Augmin complex. The x-axis represents the Pearson correlation coefficient (r), the y-axis shows the significance. Genes with significant co-dependency (q < 0.05) are shown in dark gray

**Fig. S7.**
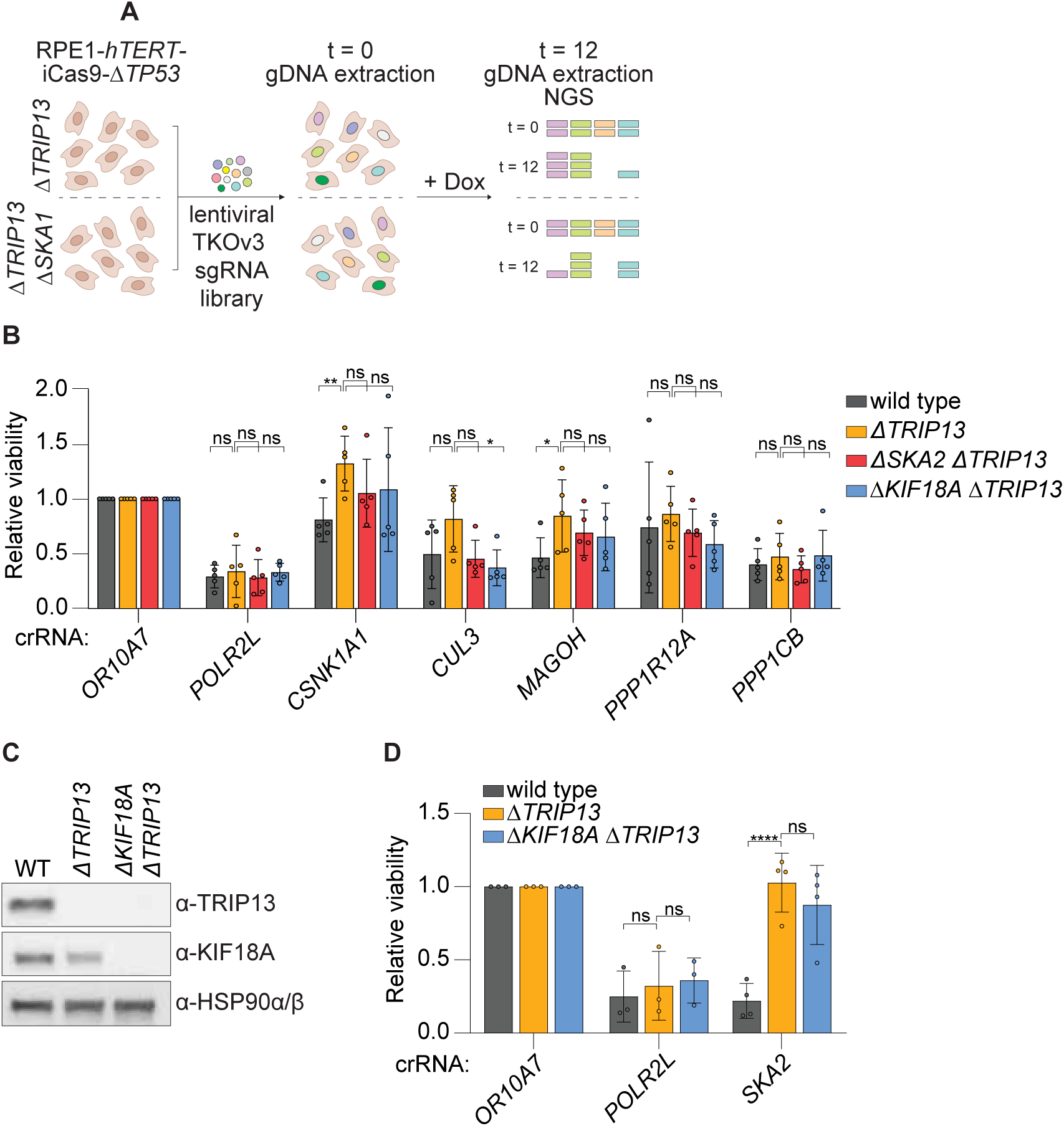
(**A**) Schematic overview of the genome-wide Δ*TRIP13* versus Δ*SKA1*Δ*TRIP13* CRISPR-Cas9 knockout screen workflow performed in triplicate. Dox, doxycycline; NGS, next-generation sequencing. (**B**) Relative viability of RPE1-*hTERT*-iCas9 Δ*TP53* (WT), RPE1-*hTERT*-iCas9 Δ*TRIP13*Δ*TP53* (Δ*TRIP13*), RPE1-hTERT-iCas9 Δ*SKA2*Δ*TRIP13*Δ*TP53 (*Δ*SKA2*Δ*TRIP13),* and RPE1-hTERT-iCas9 Δ*KIF18A*Δ*TRIP13*Δ*TP53 (*Δ*KIF18A*Δ*TRIP13)* cells upon CRISPR-based gene knockout. (**C**) Representative western blot of TRIP13 and KIF18A in WT, Δ*TRIP13*, and Δ*KIF18A*Δ*TRIP13* cells. (**D**) Relative viability upon CRISPR-based gene knockout. Data in (B) and (D) were obtained from three biologically independent experiments with technical triplicates. Statistical significance was assessed by two-way ANOVA with correction for multiple comparisons. *P < 0.05; **P < 0.01; ****P < 0.0001; ns, not significant.

**Fig. S8.**
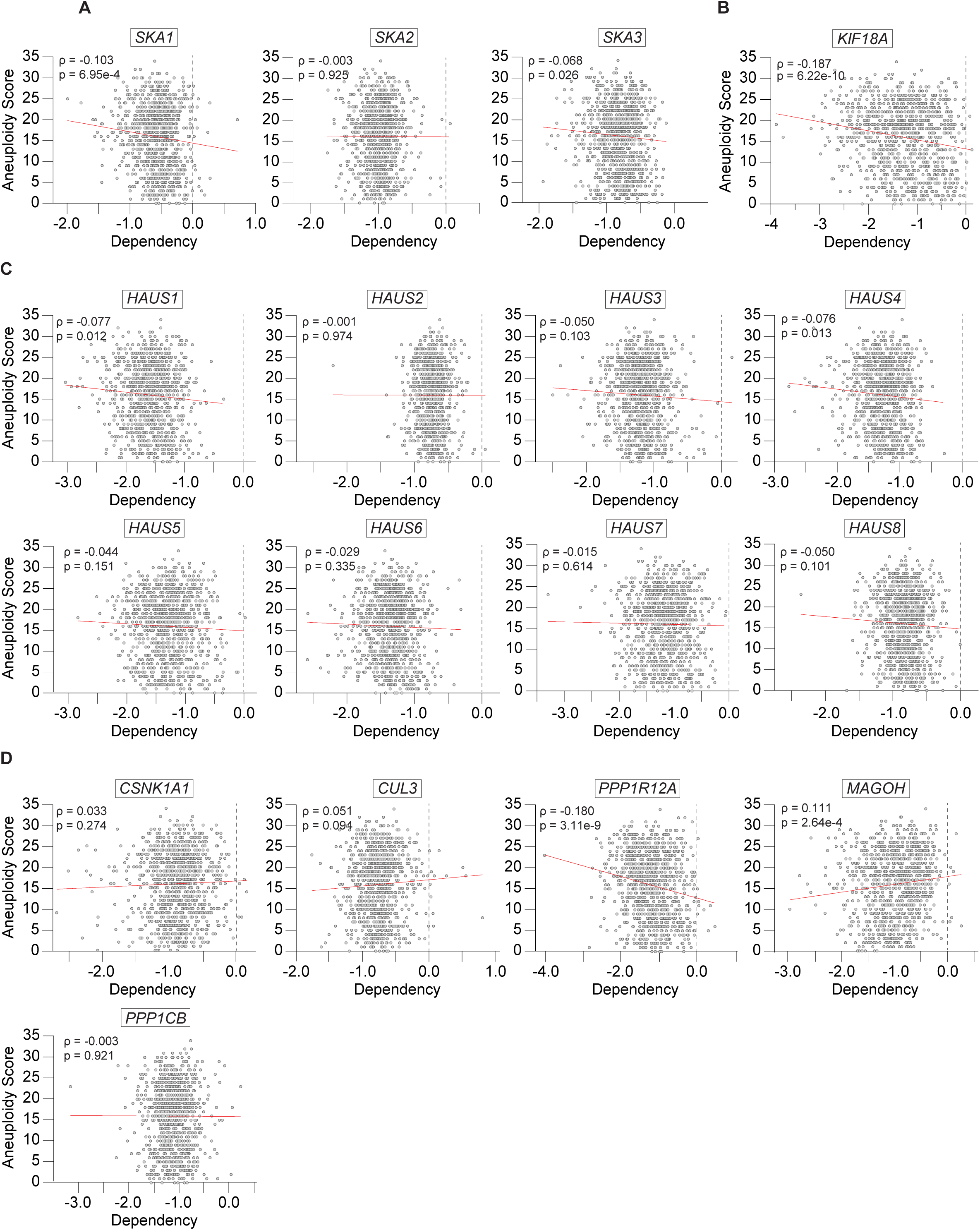
(**A**) The correlation between aneuploidy scores and the dependency on (**A**) SKA complex genes, (**B**) KIF18A, (**C**) Augmin complex genes, (**D**) mitosis-linked identified synthetic viable hits, in the Chronos CRISPR-Cas9 dataset. Pearson correlation coefficient and corresponding P value are indicated.

**Fig. S9.**
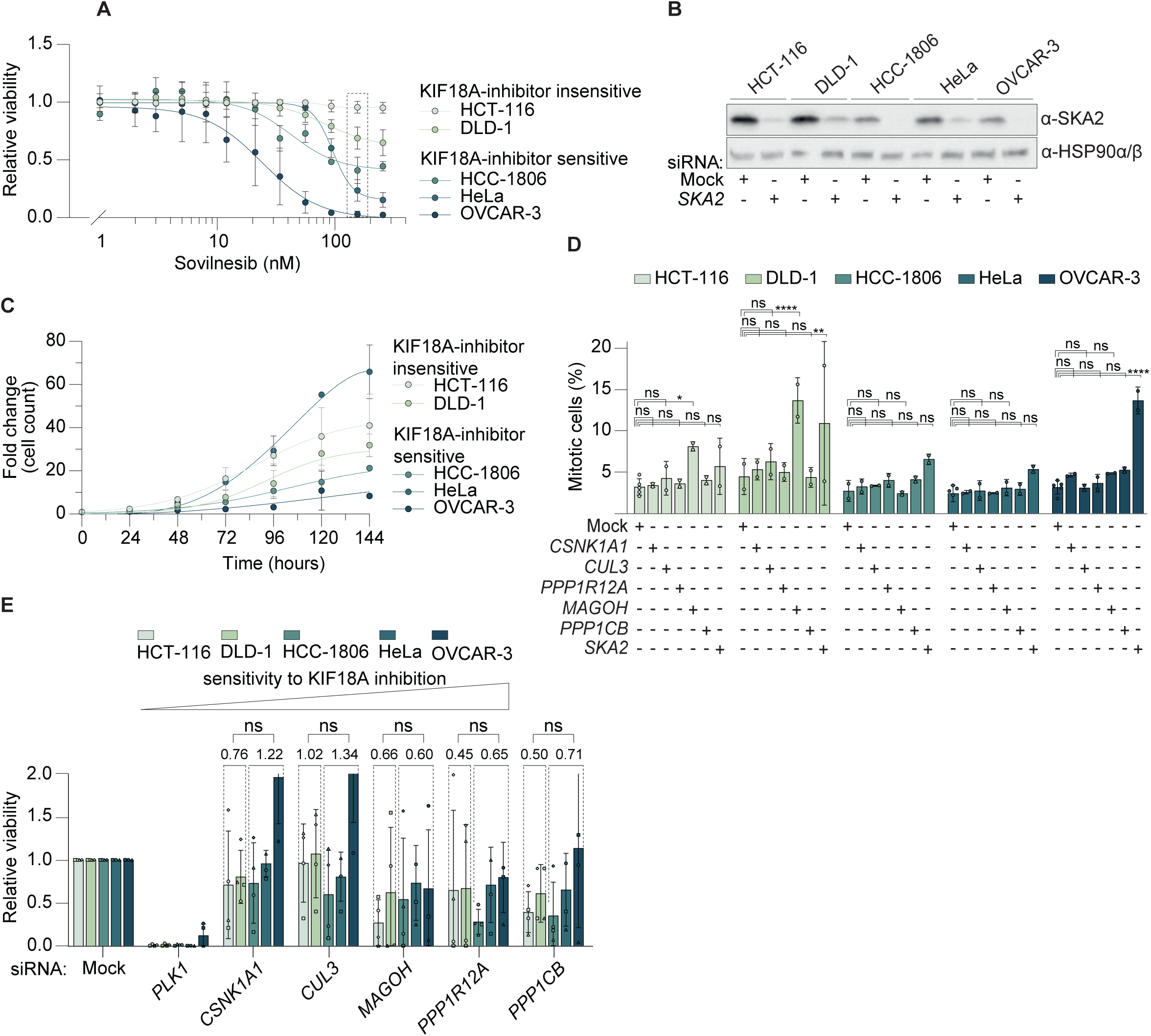
(**A**) Dose-response relative viability curve after treatment with varying concentrations of the KIF18A inhibitor Sovilnesib. Results obtained for the concentration of 155 nM, indicated with the dashed lines, are also shown in Fig. 3C. (**B**) Western blot of SKA2 levels after treatment with mock or *SKA2* siRNA. (**C**) Cell growth curve of unperturbed cancer cell lines, based on fold changes in cell counts over time. (**D**) Percentage of phospho-Histone H3-ser10 positive mitotic cells measured by flow cytometry after treatment with mock or *CSNK1A1, CUL3, PPP1R12A, MAGOH, PPP1CB,* or *SKA2* siRNA for 60 hours. (**E**) Relative viability after protein depletion, grouped as KIF18A-inhibition-insensitive (HCT-116, DLD-1) and KIF18A-inhibition-sensitive (HCC-1806, HeLa, OVCAR-3). Data were obtained in three (A) and (C), two (D), or four (E) biologically independent experiments with technical triplicates (A), (C), (E) or duplicates (D). Statistical significance was assessed by two-way ANOVA with correction for multiple comparisons. *P < 0.05; **P < 0.01; ****P < 0.0001; ns, not significant.

**Fig. S10.**
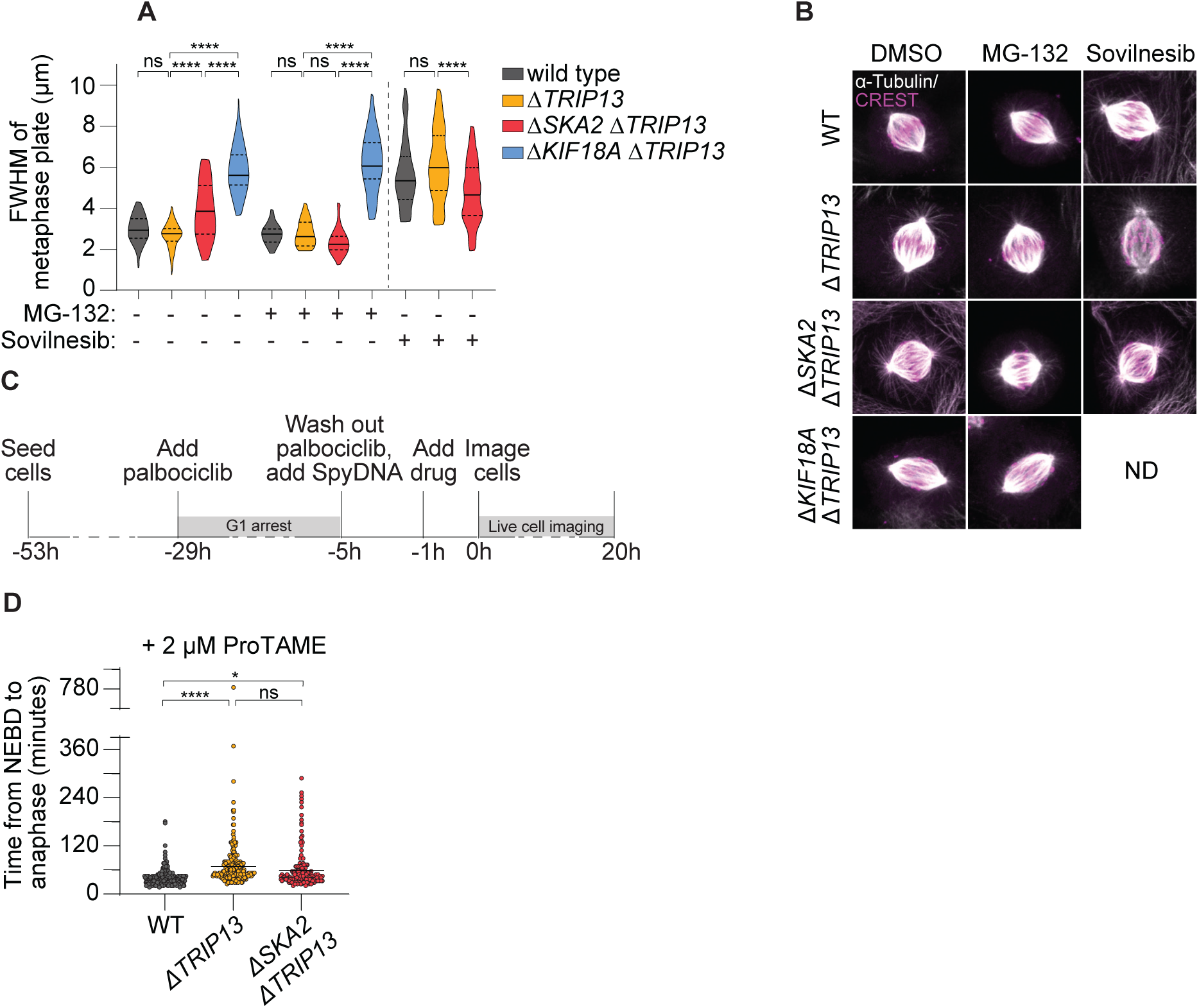
(**A**) Full width at half maximum (FWHM) of the metaphase plate of RPE1-*hTERT*-iCas9-Δ*TP53* (WT), RPE1-*hTERT*-iCas9 Δ*TP53*Δ*TRIP13* (Δ*TRIP13*), RPE1-*hTERT*-iCas9 Δ*TP53*Δ*SKA2*Δ*TRIP13* (Δ*SKA2*Δ*TRIP13*), and RPE1-*hTERT*-iCas9 Δ*TP53*Δ*KIF18A*Δ*TRIP13* (Δ*KIF18A*Δ*TRIP13*) cells, with or without MG-132 or Sovilnesib treatment. For WT, Δ*TRIP13*, Δ*SKA2*Δ*TRIP13*, and Δ*KIF18A*Δ*TRIP13*, 48, 50, 40, and 48 cells were analyzed, respectively. For MG-132-treated WT, Δ*TRIP13*, Δ*SKA2*Δ*TRIP13*, and Δ*KIF18A*Δ*TRIP13*, 45, 30, 38, and 46 cells were analyzed, respectively. For Sovilnesib-treated WT, Δ*TRIP13*, and Δ*SKA2*Δ*TRIP13,* 52, 49, and 36 cells were analyzed, respectively. (**B**) Representative images of WT, Δ*TRIP13*, Δ*SKA2*Δ*TRIP13*, and Δ*KIF18A*Δ*TRIP13* cells treated with 5 μM MG-132 or 250 nM Sovilnesib, treatments used in Fig. 4BC-D and Fig. S10B. (**C**) Schematic overview of the live-cell imaging workflow used to image the first mitosis after ProTAME treatment. (**D**) Mitotic duration (measured as time from NEBD to anaphase) of WT, Δ*TRIP13*, and Δ*SKA2*Δ*TRIP13* cells treated with 2 µM ProTAME. Average duration per condition: WT = 40 minutes, Δ*TRIP13* = 68 minutes, Δ*SKA2*Δ*TRIP13* = 59 minutes. Only cells observed entering and exiting mitosis were included in this analysis. 194, 281, and 186 cells were analyzed for WT, Δ*TRIP13*, and Δ*SKA2*Δ*TRIP13* cells, respectively. Data in (A) were obtained in three or four biologically independent experiments, data in (C) were obtained in three biologically independent experiments. Statistical significance was assessed by two-way ANOVA (A) and one-way ANOVA (C), each with correction for multiple comparisons. *P = 0.03; ****P < 0.0001; ns, not significant.

**Table S1.**
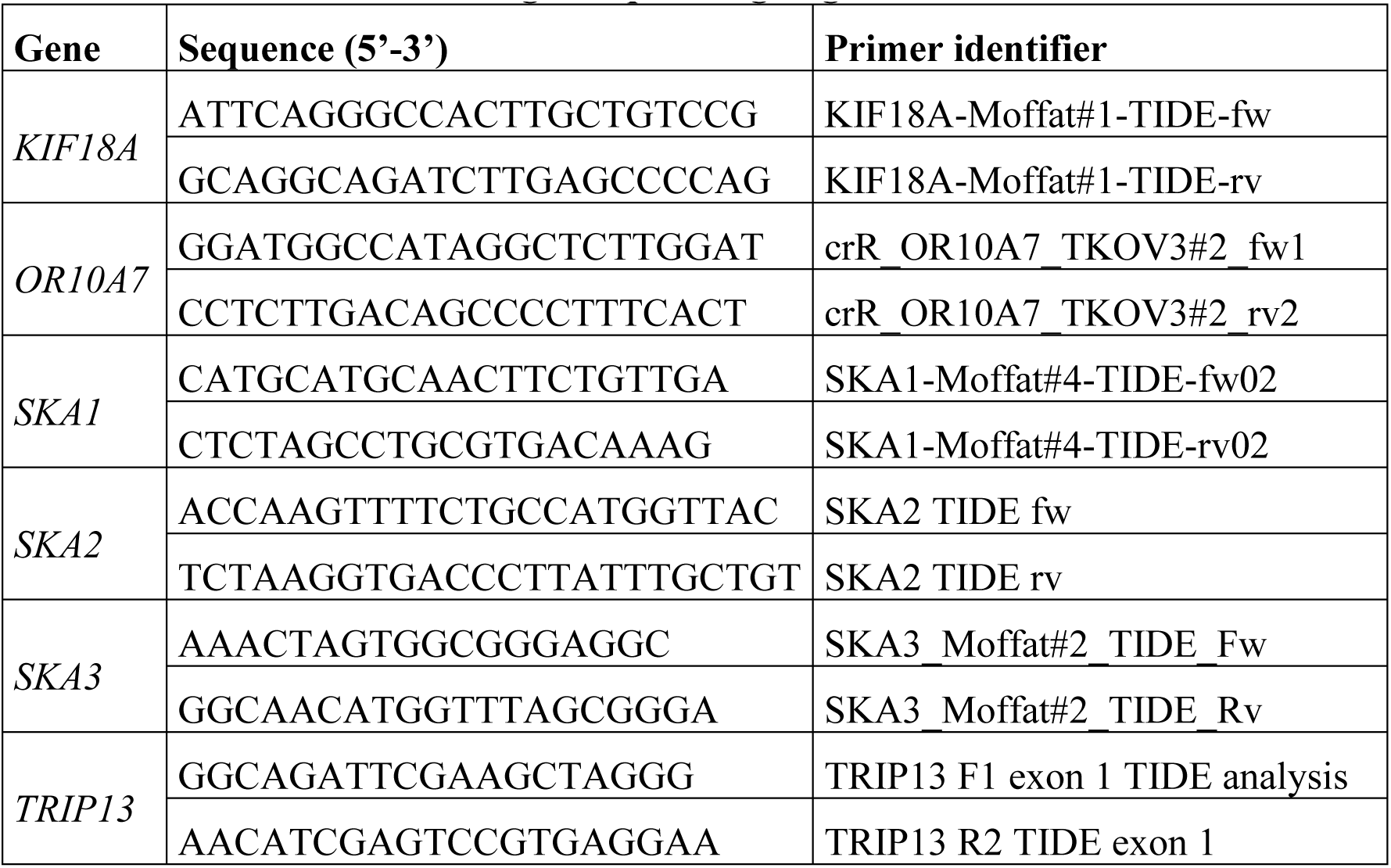
Primers used for Sanger sequencing of gene knockouts.

**Table S2.**
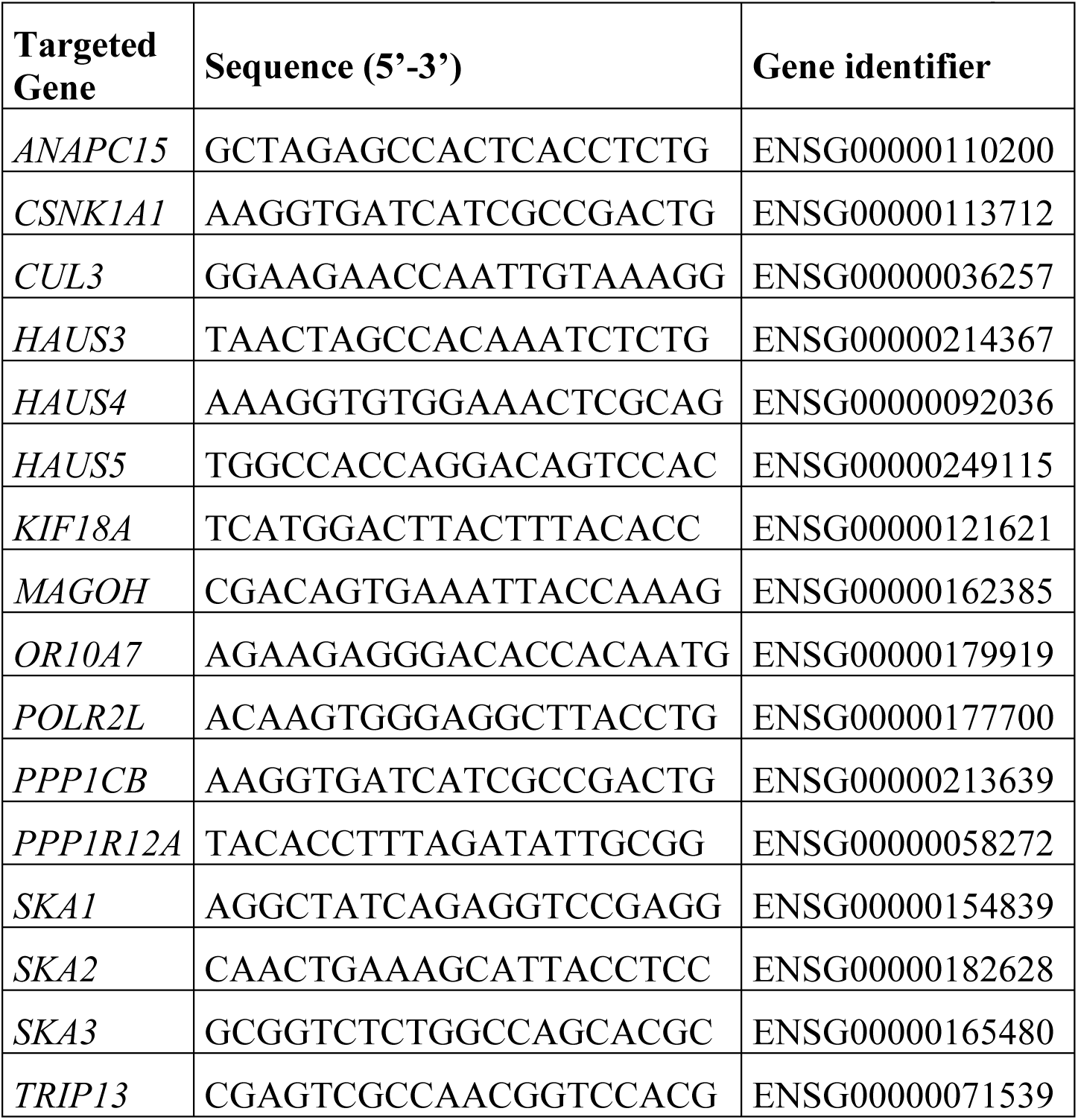
crRNA sequences used for CRISPR-Cas9 mediated genome editing.

